# Cell surface receptor kinase FERONIA linked to nutrient sensor TORC signaling controls root hair growth at low temperature linked to low nitrate in *Arabidopsis thaliana*

**DOI:** 10.1101/2022.01.10.475584

**Authors:** Javier Martínez Pacheco, Limei Song, Lenka Kuběnová, Miroslav Ovečka, Victoria Berdion Gabarain, Juan Manuel Peralta, Tomás Urzúa Lehuedé, Miguel Angel Ibeas, Martiniano M. Ricardi, Sirui Zhu, Yanan Shen, Mikhail Schepetilnikov, Lyubov A Ryabova, José M. Alvarez, Rodrigo A. Gutierrez, Guido Grossman, Jozef Šamaj, Feng Yu, José M. Estevez

**Author notes:** Co-first authors (Javier Martínez Pacheco, Limei Song). Correspondence should be addressed. (F.Y) and (J.M.E).

## Abstract

Root hairs (RH) are excellent model systems for studying cell size and polarity since they elongate several hundred-fold their original size. Their tip growth is determined both by intrinsic and environmental signals. Although nutrient availability and temperature are key factors for a sustained plant growth, the molecular mechanisms underlying their sensing and downstream signaling pathways remain unclear. Here, we identified that low temperature (10°C) triggers a strong RH elongation response involving the cell surface receptor kinase FERONIA (FER) and the nutrient sensing TOR Complex 1 (TORC). In this study, we found that FER is required to perceive limited nutrient availability caused by low temperature. FER interacts with and activates TORC downstream components to trigger RH growth. In addition, the small GTPase Rho-related protein from plants 2 (ROP2) is also involved in this RH growth response linking FER and TOR. We also found that limited nitrogen nutrient availability can mimic the RH growth response at 10°C in a NRT1.1-dependent manner. These results uncover a molecular mechanism by which a central hub composed by FER-ROP2-TORC is involved in the control of RH elongation under low temperature and nitrogen deficiency.

## Introduction

Root hairs (RH) are cell outgrowths that develop as cylindrical protrusions from the root epidermis in a developmentally regulated manner^1^. RHs are able to expand in a polar manner several hundred times their original size in a couple of hours to reach water-soluble nutrients in the soil, to promote interactions with the local microbiome, and to support the anchoring of the plant^2^. RH growth is controlled by the coordination of a plethora of environmental and endogenous factors^3,4^. Recently, an autocrine mechanism of RH growth was described where RALF1-FER complex recruits and phosphorylates the early translation initiation factor 4E1 (elF4E1) to enhance protein synthesis of specific mRNAs, including the RH growth master regulator *ROOT HAIR DEFECTIVE SIX-LIKE4 (RSL4*)^5^, The observation that RH cells can respond to their local environment in a cell-autonomous manner within minutes^6^, points towards mechanisms that directly modulate the growth machinery at the growing apex.

It is known that macronutrient availability is a key factor that promotes rapid RH growth^3,7,8^ Recently, we showed that under low temperature conditions (10°C), nutrient availability in the media is reduced and triggers an enhanced RH growth that is suppressed if nutrients are increased^9^. We specifically showed that RH growth of WT Col-0 plants is highly responsive to increasing nutrient concentrations (from 0.5X Murashige and Skoog (MS) to 2.0X MS) with regular concentration gelling agents (0.8%). High nutrient concentration impairs RH growth even if they are exposed to low temperatures. In a similar manner, an increase in agar concentration (from 0.8% to 2.5%) in the MS medium, which likely restrains nutrient mobility and nutrient uptake^10–13^ blocked low temperature-induced RH elongation^9^. These observations suggested that low temperature restrict nutrient mobility and availability in the culture medium, leading to promotion of polar RH growth. It is still unclear which specific nutrients are affected and the signaling pathways involved in the temperature effect that triggers this RH growth response. Diminished nutrient availability is known to activate RH expansion through a transcriptional reprogramming governed by the transcription factors (TF) RHD6-RSL4^9^. Specifically, a novel ribonucleoprotein complex composed of lncRNA AUXIN-REGULATED PROMOTER LOOP (APOLO) and the TF WRKY42 forms a regulatory hub to activate RHD6 by shaping its epigenetic environment and integrate low temperature signals governing RH growth^9,14^. In addition, we recently identified new molecular components involved in low temperature RH growth. We found that cell wall-apoplastic related peroxidases, PRX62 and PRX69, are important for low temperature triggered RH growth by inducing changes in ROS homeostasis and cell wall EXTENSIN insolubilization^15^. Although relevant advances have been achieved in our understanding on how RH growth occurs at low temperature, it is still unclear how nutrient availability caused by low temperature is perceived in the RH cells and the extent of molecular responses that control RH growth.

TARGET OF RAPAMYCIN (TOR) is an evolutionarily conserved Ser/Thr protein kinase in all eukaryotic organisms that acts as a central growth regulator controlling metabolism and protein synthesis^16–18^. TOR is found in at least two distinct multiprotein complexes called TOR Complex 1 and 2 (TORC1 and TORC2) in animal cells, although only the equivalent for TORC1 has been experimentally validated in plants^19,20^. The *Arabidopsis* TOR complex is encoded by one *TOR* gene (*At*TOR)^21^, two *REGULATORY-ASSOCIATED PROTEIN OF TOR (RAPTOR 1A and 1B*) genes^22–27^, and two *LETHAL WITH SEC THIRTEEN 8 (LSTB*) genes^28^. In contrast to the embryo-lethal tor-null mutant lines, *raptor1b* and *lst8-1* mutants are viable but show significant growth defects and developmental phenotypes^29^. Some canonical downstream targets of TOR are conserved in plants, such as the 40S ribosomal S6 kinase (S6K) which stimulates protein synthesis^24,30–33^. In plants, TOR complex plays a key role during many stages of the plant life cycle by controlling both anabolic and catabolic downstream processes. In addition, TOR is activated by nutrient availability and inactivated by stresses that alter cellular homeostasis^18,34–38^. The TOR complex senses and integrates signals from the environment to coordinate developmental and metabolic processes including hormones (e.g., auxin), several nutrients (e.g., nitrogen and sulfur), amino acids and glucose^16,27,39–43^. However, the underlying molecular mechanism by which TOR operates at a single plant cell level has yet to be elucidated.

To date, few upstream regulators of TOR have been described in plants. Among them, there is a subfamily of small GTPase Rho-related protein from plants (ROPs) involved in the spatial control of cellular processes by signaling to the cytoskeleton and vesicular trafficking. Particularly, ROP2 activates TOR in response to auxin and nitrogen signals^43–45^. ROP2 is described as a monomeric GTP-binding protein that participates in many cellular signaling processes, including the polar growth of pollen tubes and root hairs^46–49^. ROP activation is regulated by ROP guanine nucleotide exchange factors (ROP-GEFs) which interact with several receptor-like kinases (RLKs) including the CrRLK family member FERONIA (FER)^47^. During the elongation of RH, ROP2 is activated by the ROPGEF1-FER interaction. This process is also regulated by ROP-GEF4 and ROP-GEF10^47,50^. FER was also linked to carbon/nitrogen balance during plant growth^51^. Even more, it was shown that the cytoplasmic kinase domain of FER and its partner RIPK (RPM1-induced protein kinase) interact with TOR and RAPTOR-1B forming a complex to positively modulate the TOR pathway under low nitrogen nutrient conditions^52^. Currently, it is clear that plant growth can be regulated via the FER-TOR pathway as it links amino acid and/or nutrients signaling with true leaves development^51^. This evidence suggests that nutrient-mediated sensing at the cell surface level triggers a response using the TOR signaling pathway. However, it is still unknown which are the environmental signals activated in RH by the FER-TOR pathway in RH. Considering that low temperature stress can result in enhanced RH growth^9,14^, this stress condition was used as a proxy to investigate environmental signals that activate RH cell elongation. Our research reveals a novel mechanism in which FER, ROP2 and TOR are necessary to regulate RH growth in *Arabidopsis* in the context of the nitrogen nutrient availability as caused by low temperature stress.

## Results

### FER is required to trigger a strong RH growth response at low temperature

We asked which might be the RH surface protein involved in perceiving or transducing the low temperature stimulus. Since CrRLK1L FER was shown as an important hub between the cell surface signaling and downstream processes during RH growth^5,47^, we tested whether *fer-4* null mutant or *fer-5* (a truncated version with shortened kinase domain (KD)) can respond to low temperature stimulus **(Figure S1A)**. Both *fer* mutants failed to trigger RH growth at low temperature **(Figure 1A)**. We considered plant genotype (Geno), treatment (Temp), and the interaction between genotype and treatment (Geno:Temp) as the factors for two-way analysis of variance (ANOVA) to define significant changes in RH growth in response to cold. We determined that the genotype factor has a substantial and significant effect on RH growth in *fer-4* and *fer-5* mutants. Notably, the Geno:Temp interaction term is also significant for *fer-4* and *fer-5*. These results suggest that FER contributes to general RH growth and RH response to cold **(Figure S1B)**. In contrast, the mutant *eru*, impaired in the related CrRLK1L ERULUS (ERU), previously linked to RH growth and cell wall integrity processes with very short RH phenotype at 22°C^53,54^, was able to react to low temperature which triggered RH growth response although to a lower extent than wild-type Col-0 plants **(Figure 1A)**. This indicates that FER, but not ERU, might be involved specifically in this growth response despite being phylogenetically related^54^. Then, a fluorescent translational reporter line of FER (*p*FER:FER-GFP), was used to study *FER* expression in roots and RH. FER protein levels are increased clearly activated in root cells under low temperature stimulus (**Figure 1B)**, and during RH growth **(Figure 1C)**. Distribution of FER-GFP in the clear zone and punctate compartments in the cytoplasm was not affected by the temperature change from 22°C to 10°C. However, accumulation of FER-GFP at the plasma membrane was increased at low temperature (at 10°C) **(Figure 1C)**. Finally, phosphorylated, and non-phosphorylated levels of FER (FER-p/FER) were quantified at both temperatures. Under low temperature, almost half of the FER protein levels were present as a non-phosphorylated form and after 3 days of growth, almost all FER protein was present as a phosphorylated version **(Figure 1D)**. This indicates that low temperature not only increases FER protein levels at the plasma membrane of growing RH, but also promotes a complete phosphorylation of FER protein. This is possibly enhancing the interaction with putative partners and triggering the activation of downstream signaling components of RH polar growth.

**Figure 1.**
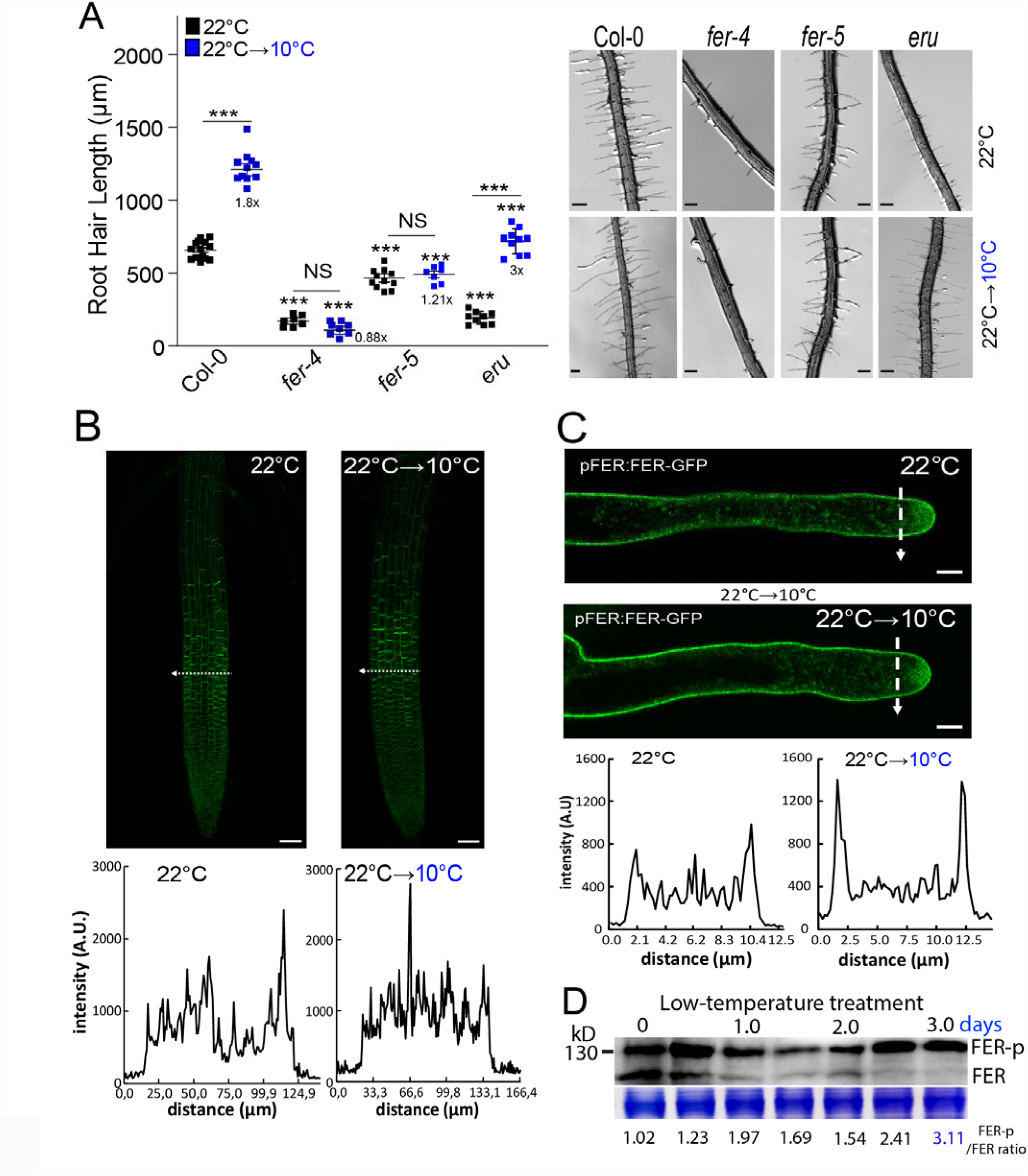
High levels of FER in its phosphorylated form are required to trigger the low temperature RH growth. **(A)** Scatterplot of RH length of Col-0, of *fer-4, fer-5*, and *eru* mutants grown at 22°C or at 10°C. RH growth is enhanced at low temperature in Wt Col-0 and *eru* mutant but not in the *fer-4* and *fer-5* mutants. Each point is the mean of the length of the 10 longest RHs identified in the maturation zone of a single root. Data are the mean ± SD (N=l0 roots), two-way ANOVA followed by a Tukey–Kramer test; (***) *p* <0.001, NS= non-significant. Results are representative of three independent experiments. Asterisks indicate significant differences between Col-0 and the corresponding genotype at the same temperature or between the same genotype at different temperatures. Representative images of each genotype are shown on the right. Scale bars= 300 μm. Numbers under the plots represents RH growth ratio 10°C/22°C. **(B-C)** Increased level of FER in root cells after its activation by low temperature. **(B)** Confocal images of root apex of a fluorescent translational reporter line of FER (*pFER:FER-GFP*) at 22°C and after transfer from ambient to low temperature (22°C→10°C). On the Bottom: semi-quantitative evaluation of the GFP fluorescence intensity across the root tip at 22°C and after transfer from ambient to low temperature at the transition zone, indicated by a interrupted white line. Fluorescence intensity is expressed in arbitrary units (AU), N=4-6 roots. Results are representative of three independent experiments. Scale bars: 50 μm. **(C)** Confocal images of short root hairs of a fluorescent translational reporter line of FER (*p*FER:FER-GFP) at 22°C and after transfer from ambient to low temperature (22°C→10°C) On the bottom: semi-quantitative evaluation of the GFP fluorescence intensity across the subapical root hair zone at 22°C and after transfer from ambient to low temperature, indicated by a interrupted white line. Fluorescence intensity is expressed in arbitrary units (A.U.), N=6-8 roots, 8-11 root hairs. Results are representative of three independent experiments. Scale bars=5 μm. **(D)** Phosphorylation levels on FER (FER-p in *p*FER:FER-GFP) increases after 3 days at low temperature in roots. Protein loading control (Coomasie Blue) is indicated below. FER-p/FER ratios were analyzed by ImageJ.

### TOR pathway is involved in low temperature induced RH growth

Recently, it was shown that FER-RIPK interacts with TOR-RAPTOR1B and phosphorylates it leading to TOR pathway activation in the context of nutrient perception and regulation of global metabolism^52^. We then asked whether TOR might be also involved in the RH growth process under low temperature. We used a β-estradiol (es)-induced TOR knockdown (*tor-es*, es-induced RNAi silencing of TOR) line^55^ which showed short RH as reported previously^32^. Notably, induction of TOR silencing with estradiol completely blocked the RH response to low temperature **(Figure 2A)**. Similarly, inhibition of TOR kinase activity with AZD-8055^56^, an ATP-competitive inhibitor of mTOR kinase activity, abolished the RH growth at both 22°C and low temperature **(Figure 2B)**. Overexpression of TOR enhanced RH growth at both temperatures assayed **(Figure 2C)**. Using two-way ANOVA analysis, we found that alteration of TOR levels impacts general RH growth (Geno factor, **Figure S2A)**. However, the significant interaction between genotype and treatment indicates TOR is also involved in RH responses to cold (Geno:Temp, **Figure S2A)**. These results suggest that the TOR pathway is involved in the RH growth and specifically in RH low temperature growth response and might operate downstream of FER. At the transcriptional level, TOR root expression level is upregulated up to three-folds at 10°C vs 22°C **(Figure S2B)**. We tested whether the growth response at low temperature in RH is also affected in mutants of the TOR pathway and some downstream components. All the plant mutants tested (*raptor1b, rps6b*, and *lst8-1*) were unable to respond to the low temperature treatment. They exhibited a broad spectrum of RH phenotypes as compared to Wt Col-0 plants **(Figure 2C)**. For instance, *raptor1b* behaved similarly to *tor-es* mutant, while *rps6b* showed an intermediate phenotype and *lst8* was comparable to Wt Col-0 plants (at 22°C) **(Figure 2C)**. In addition, overexpression of TOR and the S6 KINASE 1 (S6K1), a direct downstream target of TOR, showed enhanced RH growth at 22°C. Since previous reports showed phosphorylation of the S6K1 can be used to monitor TOR protein kinase activity in plants^32,55^, we measured ratios of S6K-p/S6K. Strong induction of the S6K-p/S6K ratio was observed in Wt Col-0 under low temperature **(Figure 2D)**. These results suggest TOR and some downstream components (e.g. S6K and RPS6b) are required to promote RH growth under low temperature.

**Figure 2.**
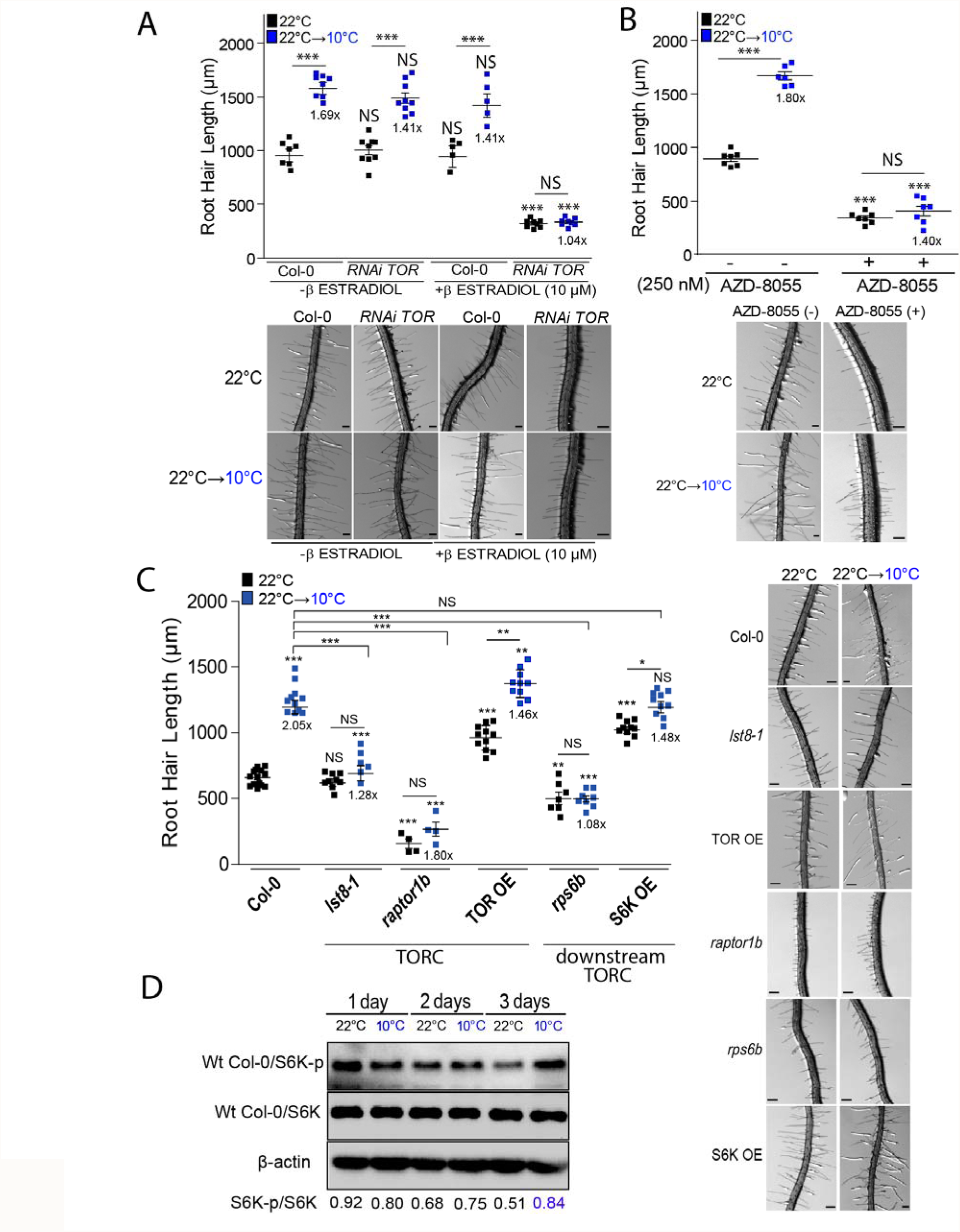
TORC signaling pathway is required for low temperature triggered RH growth. **(A)** Scatterplot of RH length of Col-0 and *tor-es* line grown at 22°C or at l0°C. Differential growth of RH at low temperature is suppressed in the estradiol inducible *RNAi TOR* line. Each point is the mean of the length of the 10 longest RHs identified in the maturation zone of a single root. Data are the mean ± SD (N=7-10 roots), two-way ANOVA followed by a Tukey–Kramer test; (***) *p*<0.001, NS=non-significant. Results are representative of three independent experiments. Asterisks indicate significant differences between Col-0 and the corresponding genotype at the same temperature or between the same genotype at different temperatures. Representative images of each line are shown below. Scale bars=300 μm. Numbers under the plots represents RH growth ratio 10°C/22°C. **(B)** Differential growth of RH at low temperature is abolished in the Col-0 treated with 250 nM of TOR inhibitor, AZD-8055. Each point is the mean of the length of the 10 longest RHs identified in the maturation zone of a single root. Data are the mean ± SD (N=7 roots), two-way ANOVA followed by a Tukey–Kramer test; (***) *p*<0.001, NS=non-significant. Results are representative of three independent experiments. Asterisks indicate significant differences between Col-0 and the corresponding genotype at the same temperature or between the same genotype at different temperatures. Representative images of each line are shown below. Scale bars=300 μm. Numbers under the plots represents RH growth ratio 10°C/22°C. **(C)** RH elongation of TOR and downstream TOR signaling pathway mutants under low temperature. Each point is the mean of the length of the 10 longest RHs identified in the maturation zone of a single root. Data are the mean ± SD (N=7-12 roots), two-way ANOVA followed by a Tukey–Kramer test; (*) *p*<0.05, (**) *p*<0.01, (***) *p*<0.001, NS=non-significant. Results are representative of three independent experiments. Asterisks indicate significant differences between Col-0 and the corresponding genotype at the same temperature or between the same genotype at different temperatures. Representative images of each line are shown on the right. Scale bars=300 μm. Numbers under the plots represents RH growth ratio 10°C/22°C. **(D)** Analysis of the phosphorylation state of S6K (S6K-p/S6K ratio) in Col-0. A representative immunoblot is shown of two biological replicates (see **Supplementary Material 1)**. S6K-p/S6K ratio was analyzed by ImageJ.

### FER facilitates TOR polar localization in RH and activation of TOR pathway at low temperature

Since TOR activation under a plethora of stimuli usually triggers phosphorylation of the downstream factor S6K (S6K-p), we asked whether low temperature responses regulated by FER might control this output. We investigated the molecular mechanisms by which TOR-S6K are induced at 10°C and we tested whether this might be mediated by FER. Since *fer-4* and *tor-es* showed similar RH phenotypes at low temperature **(Figure 2A-B)**, possibly indicating that they may act in the same pathway, we first tested whether FER and TOR interaction is enhanced under 10°C vs 22°C. By performing a co-immunoprecipitation (Co-IP) analysis, we found that FER-TOR interaction is enhanced under low temperature **(Figure 3A)**. This result suggests an interaction between FER and TOR kinase is implicated in RH growth under low temperature. This corroborates previous findings that showed direct interaction of the FER kinase domain and the N-terminal domain of TOR (aa 1-1449 of TOR, NTOR)^52^. Next, we evaluated TOR levels in growing RH by immunolocalization. We found that FER is required for the apical accumulation of TOR since TOR lost the polar pattern in *fer-4* root hairs. However, the level of RH tip-localized TOR detected was similar at both, low temperature, and control conditions (22°C) **(Figure 3B-C** and **Figure S3)**. This result indicates that localization of TOR at RH tip is dependent on FER. Then, we tested whether FER is involved in S6K activation by TOR. We measured S6K-p/S6K protein ratio under both growth conditions in Wt Col-0, *fer-4, fer-5*, and FER^K565R^ (which has been shown to abolish FER autophosphorylation and transphosphorylation in an *in vitro* kinase activity assav^57,58^, although some activity might remain^59^). Interestingly, low temperature enhanced S6K-p levels after 3 days of growth in Wt Col-0 this was partially suppressed in all three *fer* mutants tested **(Figure 3D)**. This suggests that FER controls the level of TOR activation under low temperature in RH. Collectively, these results indicate that TOR localization in the RH tip and TOR activation is dependent on FER and enhanced at low temperature.

**Figure 3.**
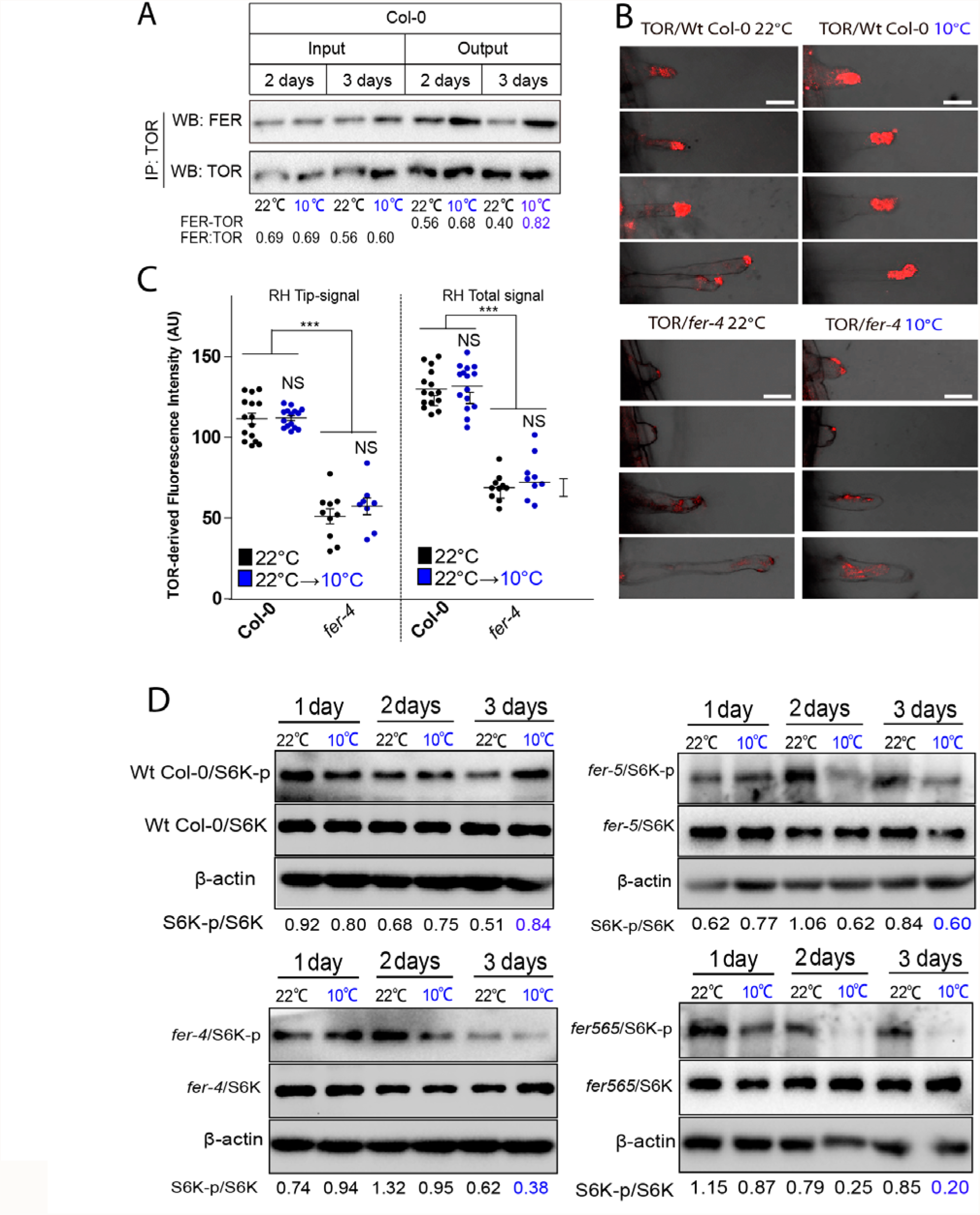
FER controls polar TOR localization and subsequent activation thus increasing phosphorylation of S6K under low temperature. **(A)** Enhanced FER-TOR interaction at low temperature in roots by lmmunoprecipitation (IP). A representative experiment of three replicates is shown (see **Supplementary Material 1)**. **(B)** Representative images showing TOR immunolocalization in RHs is FER-dependent. Scale bar=10 μm. Control images with the pre-immune serum as primary antibody are included in **Figure S3**. **(C)** Apical and total TOR signal quantification in RHs showed in (B). A ROI at the RH tip of the fluorescent signal or at the entire RH TOR-derived fluorescent signal was measured. Fluorescence AU data are the mean ± SD (N=10-15 root hairs), two-way ANOVA followed by a Tukey–Kramer test; (***) *p*<0.001, NS= non-significant. Results are representative of two independent experiments. Asterisks on the graph indicate significant differences between genotypes. **(D)** Analysis of the phosphorylation state of S6K in Wt Col-0, *fer-4, fer-5*, and *FER*^*K565R*^ mutants Arabidopsis roots. Phosphorylation of S6K (S6K-p) is enhanced at low temperature and requires FER active kinase. A representative immunoblot is shown of three biological replicates (see **Supplementary Material 1)**. S6K-p/S6K ratio was analyzed by Image J. Wt Col-0 immublot is the same of the Figure 2D.

### ROP2 is required for low temperature RH growth

Previous studies have shown a key role of ROP2 in the regulation of RH polarity and elongation^48,60,61^, as an important molecular link between FER and downstream components involved in RH growth^47^. In addition, ROP2 promotes the activation of TOR and its relocation to the cell periphery and induces the downstream signal transduction pathway^44^. More importantly, ROP2 was shown to integrate diverse nitrogen and hormone signals for TOR activation^43^. We asked whether low temperature (10°C) is able to change ROP2 targeting (as ROP2p:ROP2-mCitrine) to the plasma membrane in roots and RH. Low temperature (10°C) led to noticeable increase of ROP2 fluorescence intensity in root cells **(Figure 4A)**, in the apical and subapical cytoplasm, and at the apical plasma membrane of RH **(Figure 4B)**. Then, we observed that *rop2* and *rop2 rop4* partially abolishes the differential growth responses at 10°C and produces a very short RH phenotype **(Figure 4C)**. On the contrary, ROP2 OE triggers a constitutive RH growth at both temperatures **(Figure S4)** similarly to *S6K1 OE* and *TOR OE* lines **(Figure 2C)**. In addition, when ROP2 OE is expressed in the *fer* mutant background *fer-8* (ROP2 *OE/fer-8;* **Figure S4)**, this strong growth effect disappears highlighting the role of FER on the ROP2 function. On the other hand, while high levels of TOR enhance RH growth regardless of the temperature treatments, the constitutively active ROP2 version (CA-ROP2) is able to abolish TOR activation of RH growth **(Figure 4C)**. The levels of S6K-p/S6K in TOR OE, with constitutive long RH at both temperatures, was reduced by the presence of CA-ROP2 (in TOR OE/CA-ROP2), which correlates with a diminished RH growth **(Figure 4D)**. Taken together, we suggest a direct function of ROP2 (and possibly ROP4) at the RH tip. ROP2 function is dependent on the presence of FER and it modulates TOR activity in the RH growth responses to low temperature.

**Figure 4.**
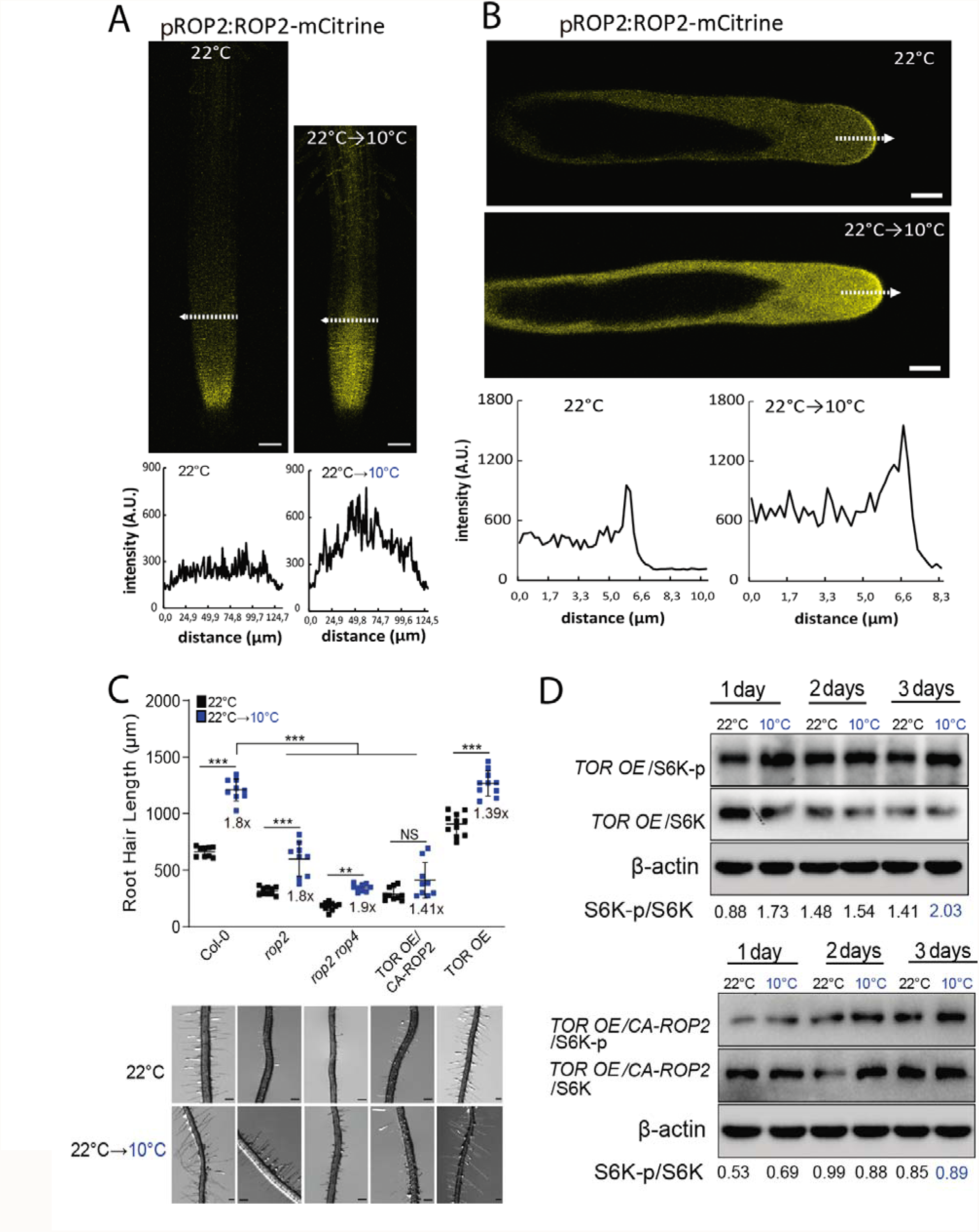
ROP2 is required to triggers RH growth at low temperature. **(A-B)** Increased level of ROP2 after its activation by low temperature. **(A)** Confocal images of root apex of a fluorescent translational reporter line of ROP2 (pROP2:ROP2-mCitrine) at 22°C (A) and after transfer from ambient to low temperature (22°C→10°C). On the bottom: semi-quantitative evaluation of the mCitrine fluorescence intensity across the root tip at 22°C and after transfer from ambient to low temperature at the transition zone, indicated by a interrupted white line. Fluorescence intensity is expressed in arbitrary units (A.U.), N=4-5 roots. Results are representative of three independent experiments. Scale bars=50 μm. **(B)** Increased level of ROP2 in the plasma membrane of RHs after its activation by low temperature. Confocal images of short RHs of a fluorescent translational reporter line of ROP2 (ROP2p:ROP2-mCitrine) at 22°C and after transfer from ambient to low temperature (22°C→10°C). On the bottom: semi-quantitative evaluation of the mCitrine fluorescence intensity across the RH apex at 22°C and after transfer from ambient to low temperature, indicated by a interrupted white line. Fluorescence intensity is expressed in arbitrary units (A.U.), N=4-5 roots, 6-8 root hairs. Results are representative of three independent experiments. Scale bars=5 μm. **(C)** ROP2 is required for RH at low temperature meanwhile *rop2, rop2 rop4* and *CA-ROP2/TOR OE* blocks RH growth. Each point is the mean of the length of the 10 longest RHs identified in a single root. Data are the mean ± SD (N=10-15 roots), two-way ANOVA followed by a Tukey–Kramer test; (**) *p*<0.01, (***) *p<0*.*001*. Results are representative of three independent experiments. Asterisks indicate significant differences. Representative images of each line are shown below. Scale bars=300 μm. Numbers under the plots represents RH growth ratio 10°C/22°C. **(D)** Analysis of the phosphorylation state of S6K in *TOR OE* and *TOR OE /CA-ROP2* Arabidopsis roots. Phosphorylation of S6K (S6K-p) is enhanced in *TOR OE* and suppressed in *TOR* OE/ROP2-CA. A representative immunoblot is shown of three biological replicates (see **Supplementary Material 1)**. S6K-p/S6K ratio was analyzed by Image J.

### Nitrate perception and transport mediated by NRT1.1 controls RH growth at low temperature

Nitrate is a key nitrogen nutrient required for plant growth and development^62–64^. The CHLORINA1/NITRATE TRANSPORTER1.1 (CHL1/NRT1.1) is the only known nitrate receptor (transporter and receptor)^65–67^ as well as for auxin transport^68^. NRT1.1 belong to the low affinity nitrate transporter family. However, when NRT1.1 is phosphorylated at threonine 101, it behaves as a high-affinity NO_3_^-^ (nitrate) transporter, giving this protein a dual-affinity capability^69–72^. Previous studies have shown that nitrogen and specifically nitrate is important for TOR signaling^43,73^. Since low temperature growth conditions reduce nutrient availability in the media and trigger a strong response in RH growth^9,14^, we hypothesized that the nitrate signaling pathway was involved. We directly test this hypothesis, and we measured using a qualitative method the temperature effect of NO_3_^-^ diffusion in agar plates at 22°C and at 10°C **(Figure S5)**. As expected, the NO_3_^-^ diffusion at 22°C was twice as high as at 10°C, confirming our previous results that the temperature effect affects the amount of nitrates available for the roots and RHs^9,14^. In agreement, we found that high levels of nitrate (18.8 and 37.6 mM) supplied in the M407 media without nitrogen (see methods) were able to partially repress low temperature-mediated RH growth while low levels of nitrate (0.5 mM) did not affect growth under this temperature **(Figure 5A)**. As a reference, previous experiments were carried out with 0.5X MS media containing 9.3 mM of nitrate and several other compounds (see **Table S1)**. In addition, an increase in agar concentration in the 407 media (from 0.8% to 1.5%) even with high NO_3_^-^ (18.8 mM), which likely restrains NO_3_^-^ diffusion mobility induced RH expansion independently of the temperature **(Figure S5)**. Three-way ANOVA analysis across all conditions **(Figure S6A and S6B)** reveals that temperature (T), the interaction of agar with NO_3_^-^ (agar:N); and the interaction of agar, temperature, and NO_3_^-^ (agar:T:N) largely explain the variance of RH elongation; while NO_3_^-^ and agar factors on their own are less influential **(Figure S5C)**. Altogether, these observations suggest that low temperatures restrict NO_3_^-^ diffusion and availability in the culture medium, leading to the promotion of polar RH growth. This result indicated nitrate can impact RH cell elongation at low temperature, particularly under low nutrient mobility environment. Next, we tested whether NRT1.1 was involved in this low temperature RH growth response. The NRT1.1 null mutants *chl1-5, chl1*.*9* (NRT1.1 harbors a substitution P492L that suppresses its root nitrate uptake activity^8^), as well as the *CHL1*^*T101D*^ (which mimics phosphorylated version of NRT1.1) showed strong RH growth response regardless of temperature conditions **(Figure 5B)**, similarly to *S6K1 OE* and *TOR OE* lines **(Figure 2C)**, but to a lower extent in terms of RH elongation. Only the *CHL1*^*T101A*^ (dephosphorylated version of NRT1.1) was able to slightly respond to the change in temperature. As expected, similar levels of S6K-p/S6K were detected in the *chl1-9* line at both temperatures **(Figure 5C)**. When these NRT1.1 mutants were grown under high (18.8 mM) or low (0.5mM) nitrate concentrations (high N/low N) at both temperatures (at 22°C and at 10°C), a similar RH phenotype was detected between both nitrate conditions either at 22°C or 10°C **(Figure 5D)**. In contrast, Wt Col-0 RH was sensitive to both low temperature and nitrate levels. Taken together, these results suggest low temperature may restrict nitrate accessibility to the RH linked to a lower mobility in the media affecting NRT1.1-mediated signaling upstream of TOR-S6K activation.

**Figure 5.**
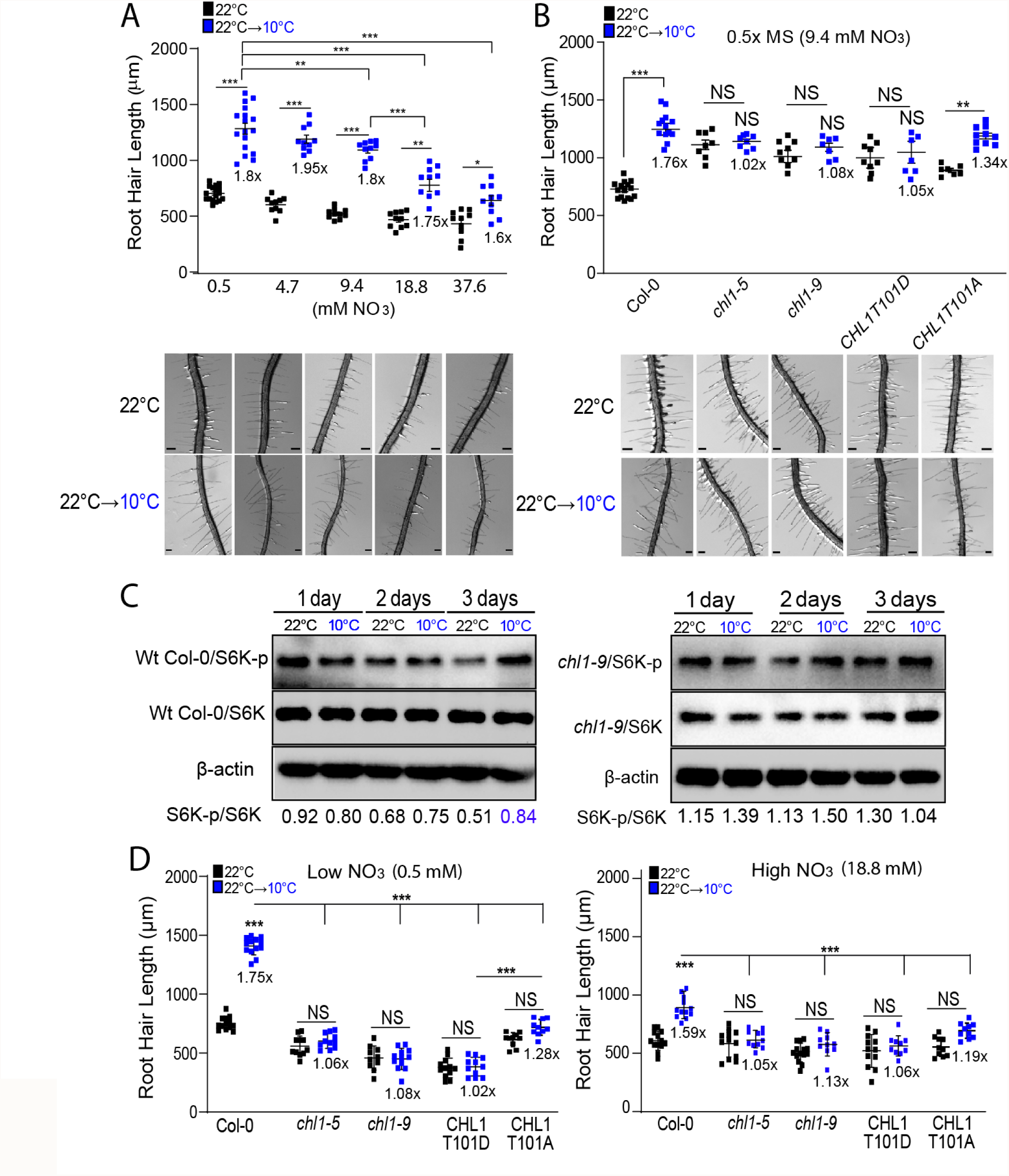
Nitrate acts as a RH growth signal perceived by NRT1.1 and triggered by TOR pathway at low temperature growth. **(A)** High nitrate supplemented in M407 media (18.8 and 37.6 mM) partially suppresses low temperature enhanced RH growth in Wt Col-0 plants. Each point is the mean of the length of the 10 longest RHs identified in the maturation zone of a single root. Data are the mean ± SD (N=10 roots), two-way ANOVA followed by a Tukey–Kramer test; (***) *p<0*.*001*. Results are representative of three independent experiments. Asterisks indicate significant differences between 0.5mM N concentration and every other N concentration at the same temperature or between the same N concentrations at different temperatures. Representative images of Col-0 under the different nitrate concentrations are shown below. Scale bars=300 μm. Numbers under the plots represents RH growth ratio 10°C/22°C. **(B)** Low temperature RH growth is regulated by the nitrate sensor NRT1.1 (CHL1). Scatter-plot of RH length of Col-0 and NRT1.1 mutants in 0.5X MS (contains 9.4 mM Nitrates) grown at 22°C or 10°C. Each point is the mean of the length of the 10 longest RHs identified in the maturation zone of a single root. Data are the mean ± SD (N=l0-15 roots), two-way ANOVA followed by a Tukey–Kramer test; (***) *p*<0.001, NS=non-significant. Results are representative of three independent experiments. Asterisks indicate significant differences between Col-0 and the corresponding genotype at the same temperature or between the same genotype at different temperatures. Representative images of each line are shown below. Scale bars=300 μm. Numbers under the plots represents RH growth ratio 10°C/22°C. **(C)** Analysis of the phosphorylation state of S6K in Wt Col-0 and *chl1-9* mutant roots. A representative immunoblot is shown of three biological replicates (see **Supplementary Material 1)**. S6K-p/S6K ratio was analyzed by ImageJ. Wt Col-0 immunoblot is the same of the Figure 2D and Figure3D. **(D)** RH growth responses under high and low nitrate (18.8 mM and 0.5 mM, respectively) in Wt Col-0, *chl1-5, chl1-9* mutants, and *CHL1^T101D^* grown at 22°C or 10°C. Each point is the mean of the length of the 10 longest RHs identified in the maturation zone of a single root. Data are the mean ± SD (N=10-15 roots), two-way ANOVA followed by a Tukey–Kramer test; (***) *p*<0.001, NS= non-significant. Results are representative of three independent experiments. Asterisks indicate significant differences between Col-0 and the corresponding genotype at the same temperature or between the same genotype at different temperatures. Numbers under the plots represents RH growth ratio 10°C/22°C.

### FER-ROP2-TOR hub is required for nitrate starvation-mediated RH growth response

Recently it was reported that RALF1-FERONIA complex is able to interact with TOR kinase under low N conditions modulating the TOR downstream signaling pathway^52^. As described above, we found that Col-0 grown at 22°C under high N condition have a shorter RH phenotype as compared to low N condition. This growth response is consistent with previous studies in which the RH length in *Arabidopsis* and other species decrease as the nitrate concentration increases although the molecular mechanism remained unknown^60,74^. To gain a better insight into the role of FER and TOR in this process, we decided to test the *fer* mutants, the TOR pathway mutants and selected overexpressing lines (TOR and S6K) in low (0.5 mM) and high nitrate (18.8 mM) media conditions **(Figure 6A** and **Figure S7)**. Under low nitrate conditions, the RH response was suppressed in both *fer* mutants and in the *tor-es* inducible mutant compared to Col-0. This result supports the idea that FER and TOR are necessary for the RH growth mediated by low nitrate conditions. The *TOR OE* line showed a RH growth phenotype similar to Col-0 under low N and a higher RH growth in excess of N. It is well known that nitrogen starvation causes enhanced root and RH growth, whereas an excess of nitrate inhibits primary root growth^75^, because of osmotic stress^71^. Furthermore, plants overexpressing TOR in a nitrate excess medium have a longer primary root phenotype^71^. According to our data that *TOR OE* line showed longer RH than Col-0 under normal conditions **(Figure 6A)**. The constitutive overexpression of TOR can lead not only to a longer primary root but also to longer RH when plants are grown on a high nitrate media to alleviate the nutrient stress. The constitutive active mutant of ROP2 suppressed the TOR OE RH phenotype while *rop2* mutant was unable to respond to contrasting levels of nitrates. A model was proposed where ROP GTPases and the cytoplasmic and apoplastic pH fluctuations can regulate RH tip growth in a nitrogen supply dependent manner thus maintaining the unidirectional growth during the RH elongation^77^. Additionally, nitrate and ammonium levels restore TOR activation under nitrogen-starvation condition via ROP2 activation^43^. We then tested if other components on the TOR complex are also involved in the low nitrate perception linked to RH growth **(Figure 6B)**. Similarly to their low temperature phenotype, *raptor1b* and *lst8-1* do not respond with differential RH growth under low nitrate. Mutation of RAPTOR1B resulted in a strong reduction of TOR kinase activity, leading to massive changes in carbon and nitrogen metabolism ^78^. The RH phenotype derived from the overexpression of S6K resembled that of TOR OE line **(Figure 6A, 6B)**. Altogether, our results show that low-nitrate (low N) RH responses are similar to those determined for low temperature. This is consistent with the idea that nitrate is one of the main signals that trigger RH expansion upon perception by NRT1.1, and subsequently transduction by FER, ROP2 and TOR pathway. We then tested how changed are the levels of S6K-p/S6K as a readout of TOR activation in low N and high N conditions in FER, TOR silenced line (*tor-es*) and NRT1.1 mutants compared to Wt Col-0 **(Figure 6C)**. First, Wt Col-0 showed an increased in S6K p/S6K ratio in low nitrogen similarly to the low temperature effect. On the contrary, and as expected, *fer-4, fer-5* and *tor-es* showed much reduced levels of S6K-p/S6K in both nitrate conditions but much lower in low nitrogen. Importantly, the effect of low N on increasing S6K-p levels is abolished in NRT1.1 mutants (*chl1-9, chl1-5* and CHLT101D), which is similar to the results observed under low temperature conditions. These results indicate that nitrate effect on S6K-p/S6K ratio depends on NRT1.1 and the FER-TOR complex.

**Figure 6.**
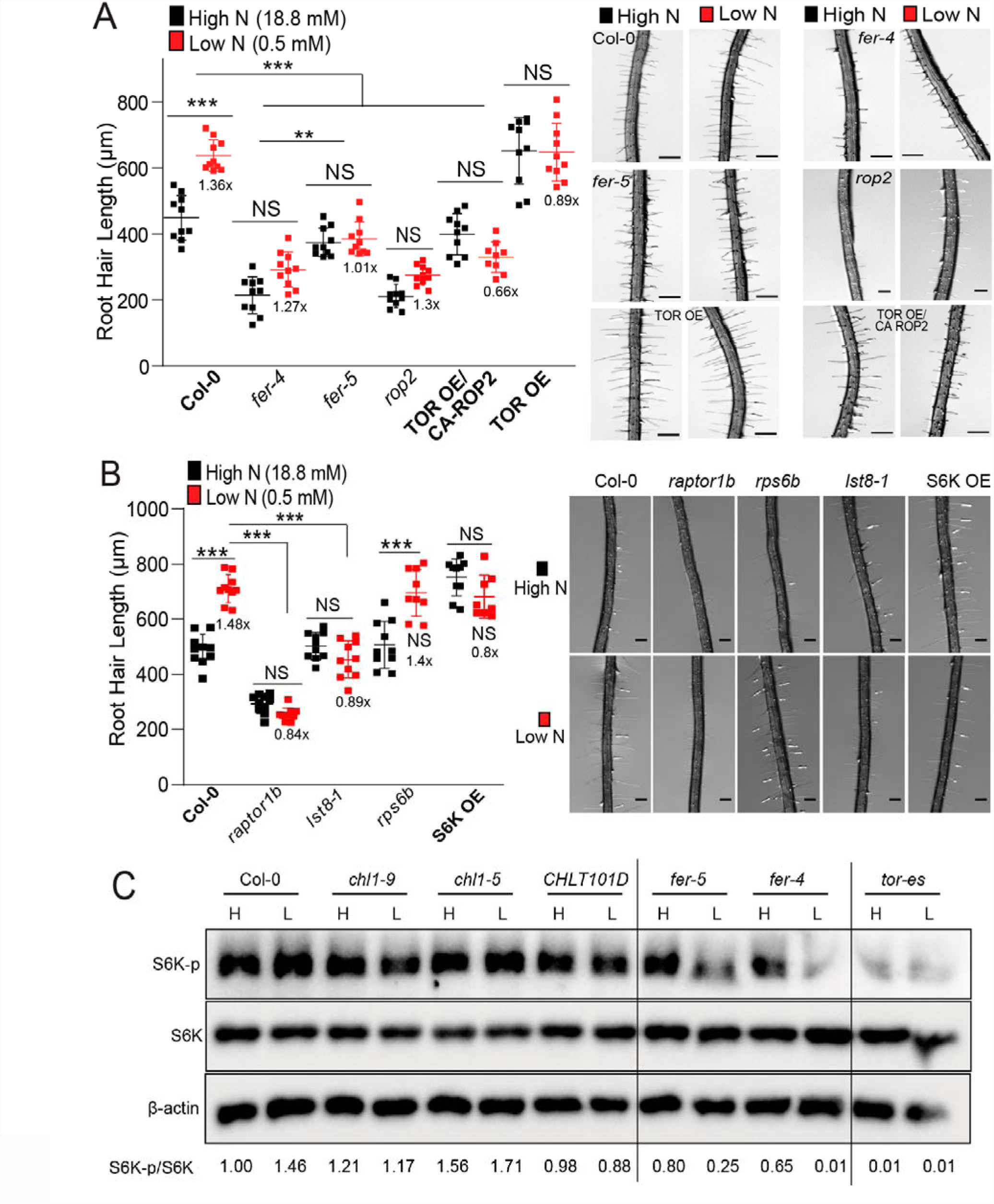
Low nitrate perception relays in FER, ROP2 and TOR to trigger RH growth. **(A)** Scatterplot of RH length of Col-0, *fer-4, fer-5, rop2*, TOR OE/CA-ROP2 and TOR OE grown in low (0.5 mM) and high nitrate (18.8 mM) conditions in M407 media, both at 22°C. Each point is the mean of the length of the 10 longest RHs identified in the maturation zone of a single root. Data are the mean ± SD (N=10 roots), two-way ANOVA followed by a Tukey–Kramer test; (**) *p*<0.01, (***) *p*<0.001, NS=non-significant. Results are representative of three independent experiments. Asterisks indicate significant differences between the same genotype at different N concentration or between different genotypes at the same N concentration. Representative images of each line are shown on the right. Scale bars=300 μm. Numbers under the plots represents RH growth ratio low N/high N. **(B)** Scatterplot of RH length of Col-0, *raptor1b, lst8-1, rps6b* and S6K OE mutants grown in low and high N conditions. Each point is the mean of the length of the 10 longest RHs identified in the maturation zone of a single root. Data are the mean ± SD (N=10 roots), two-way ANOVA followed by a Tukey–Kramer test; (***) *p*<0.001, NS=non-significant. Results are representative of three independent experiments. Asterisks indicate significant differences between the same genotype at different N concentration or between different genotypes at the same N concentration. Numbers under the plots represents RH growth ratio low N/high N. **(C)** Analysis of the phosphorylation state of S6K in Wt Col-0, in NRT1.1 mutants (*chl1-9, chl1-5*, CHLT101D), and in FER mutants (*fer-4, fer-5*, and *FER*^*K565R*^). A representative immunoblot is shown of a two Image J. biological replicates (see **Supplementary Material 1)**. S6K-p/S6K ratio was analyzed by

Since an integrative gene regulatory network analysis of TF–target interactions positioned TGA1 and its homolog TG4 as the most influential TFs in the nitrate response^79,80^, we decide to test if they play a role in the response to low temperature in RHs and low/high levels of Nitrogen **(Figure S8)**. As expected *tga1 tga4* double mutant failed to respond to low temperature and low nitrogen while TGA1 OE showed a constitutive growth response regardless the temperature and nitrogen levels. This confirms that nitrate, NRT1.1 and its downstream signaling pathway including the transcriptional responses controlled by TGA1-TG4 act an important pathway in RH growth at low temperature.

## Discussion

Growth and development of plants and animals are based on nutrient and hormonal signaling that constitute as the main regulatory networks in eukaryotes. Unraveling the functions of the regulatory hubs and their detailed molecular mechanisms are critical for deeper understanding of these signaling pathways. In all cells, there is a central hub composed by the evolutionarily conserved TOR protein kinase that integrates nutrient and energy levels coupled to stress signaling networks to further modulate cell growth^81–84^. In addition, the cell surface receptor FER acts as a versatile sensor of signals coming from the environrnent^51,85,86^. Specifically, our study uncovers a new molecular mechanism by which plants use FER-ROP2-TOR signaling pathway to control RH elongation under low nutrient conditions, specifically nitrate, induced by low temperature stress **(Figure 7)**.

**Figure 7.**
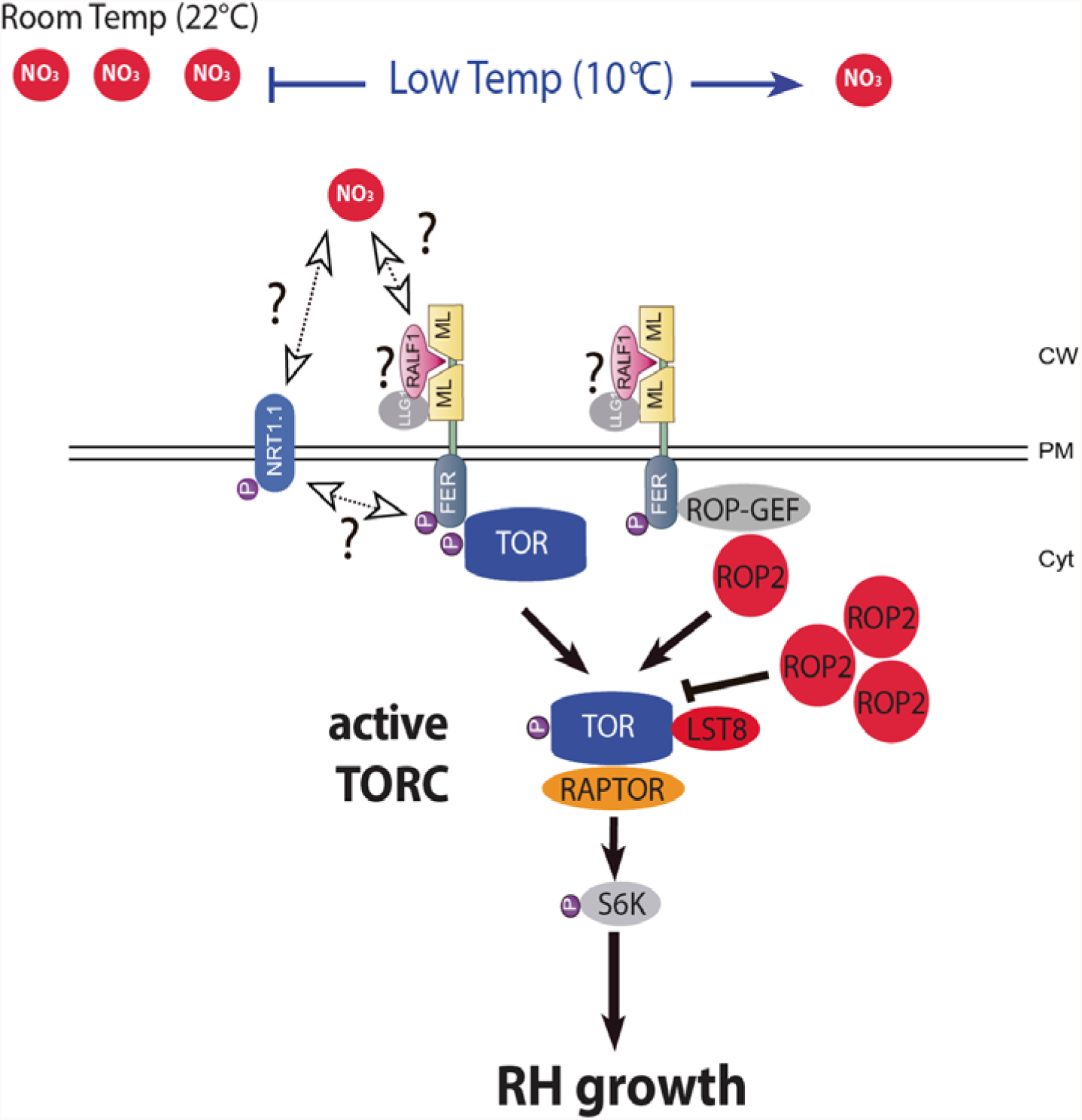
Model of FER-ROP2-TOR function in RH elongation at limited nutrient availability caused by low temperature. The model is based on the current evidence shown in this work and together with previous works. We have demonstrated that low temperature and nitrate conditions trigger RH elongation response through activation of the receptor kinase FERONIA. FERONIA triggers further activation of TOR either via direct binding to TOR (Left) or induction of the GTPase ROP2-TOR-S6K1 signaling axis via activation of GTPase ROP2 (Right). However, high levels of active ROP2 are able to negatively repress TORC activity. FERONIA can also activate NRT1.1 to promote RH elongation. Several questions (?) remain to be answered in future studies such as how low nitrate (triggered by low temperature) modulates FER-TOR activation? Does FER activate NRT1.1 directly? Do RALF1 and other RALFs respond to low nitrate to activate FER and downstream FER-TOR signaling?

In this work, we discovered that the cell surface FER receptor regulates TOR apical localization and further downstream activation, both controlling S6K phosphorylation linked to RH growth. Based on the results obtained and previous evidences, here we propose a model in which the FER-ROP2-TOR axis might be able to regulate the RH growth under low temperature with remarked attention to variable nitrate conditions **(Figure 7)**. In agreement with this, it was previously shown that ROP2 in response to auxin is an upstream effector of TOR activation^44^ and by inorganic and organic nitrogen inputs^43^. In addition, FER was shown to directly activate the TOR/RAPTOR1B signaling pathway under low inorganic and organic nitrogen conditions^52^ and is an upstream regulator of the ROPGEF4-ROP2 signaling pathway that controls ROS-mediated RH development^47,87^. In addition, ROP2 promotes TOR accumulation close to the cell periphery^44^, suggesting that plasma membrane-localized FER together with ROP2 function in regulating the relocation of TOR, close to the RH tip. Interestingly, such a local, non-genomic regulatory circuit may also explain the rapid, cell-autonomous growth regulation in RH cells observed under asymmetric conditions in the dual-flow-RootChip^6^. All these findings are in line with our results and support the idea of the existence of the FER-ROP2-TOR pathway being highly relevant for RH growth in a changing nutritional environment. Interestingly, we recently showed that FER regulates localized protein synthesis during polar RH cell growth^5,88^, suggesting that FER-TOR may also regulates polar protein synthesis in RHs. All the data together clearly highlight a key role of the FER-ROP2-TOR pathway as a core complex in sensing nutrients (e.g. nitrates) to direct RH growth.

Nitrogen is an essential nutrient for plants that when scarce limits plant growth. Although inorganic nitrogen (e.g. nitrate) is a major source of nitrogen for plants in soil, organic forms of nitrogen, like amino acids, can also be assimilated by plants from soil^89,90^. Changes in nitrate concentrations in the media mimicked this response at low temperature and NRT1.1 mutants are insensitive at RH growth level regardless the temperature or nitrate levels. We hypothesize that low nitrate in the media might rapidly induce the levels of mature-active RALF1 (and possibly other RALF peptides), which might then activate the FER-ROP2-TOR pathway and the downstream fast cell elongation effect to search for further nutrients sources. This autocrine mechanism dependent on RALF1-FER growth activation has been demonstrated for RH^88^, but not in a context that involves low temperature/low nitrate and TOR pathway. It is expected that low nitrates levels will impact on amino acid homeostasis. In line with our results, branched-chain amino acids have been shown to serve as upstream activators of TOR in plants^40^. TOR is also involved in mediating amino acid-derived metabolic regulatory signals that influence respiratory activity and plant metabolic rate^91^. Specifically, a total of 15 proteinogenic amino acids can reactivate TOR under inorganic-N starvation conditions^43^. In addition, the FER-TOR pathway responds to Gln, Asp, and Gly, reinforcing that amino acids serve as conserved upstream regulators of the TOR signaling pathway in animals and plants^52^. It is tempting to speculate that under low nitrates condition (and low temperature), the RH cell compensates with an altered amino acid homeostasis to trigger TOR activation linked fast cell growth.

Finally, several questions in our proposed model remain to be answered in future studies. How does low nitrate concentration (low temperature) triggers FER-TOR activation?. Does RALF1 and other RALFs respond to low nitrate to activate FER?. Does FER activate NRT1.1 directly?. Recently, it was shown that FER-regulated ROP2 triggers a new mechanism of negative regulation in the case of rhizosphere microorganisms such as *Pseudomonas*^76^. Although not tested here, FER ROP2-TOR signaling coupled to enhanced RH growth under low temperature (low-nutrients) might be linked to the root growth in specific soil conditions to select favorable microbiota in the soil. This pathway may integrate complex signals from soil nutrients and microbiota in the rhizosphere to plant root cell growth mechanism.

## Experimental Procedures

### Plant Material and Growth Conditions

*A. thaliana* Col-0 ecotype was used as a wild-type plant. To test the low and high nitrate response a 0.5X M407 medium without N, P nor K (M407, PhytoTechnology Laboratories, https://www.phytotechlab.com/) supplemented with 0.8% plant agar (Duchefa, Netherlands); 1.17mM MES, 0.625 mM KH_2_PO_4_ monobasic and 0.5mM KNO_3_ (low nitrate medium) 9.4mM, 18.8 mM and 37.6 mM KNO_3_ (high nitrate medium) was used. Media composition is detailed in **Table S1**. Plants were grown in the above media in continuous light (120 μmol.seg^-1^.m^-2^) either: 8 days at 22°C in the case of *tor-es* line assay, 8 days at 22°C for nitrate response assay or 5 days at 22°C + 3 days at 10°C for the low temperature and nitrate response combination assay. For the rest of the low temperature experiments plants were grown on regular 0.5X MS agar plates. For the imaging of fluorescence intensity distribution in root tips and RH, seedlings were grown for 3 days at 22°C + 3-4 days at 10°C. All mutants and transgenic lines are listed in **Table S2**. *chl1-5, chl1-9* and T101D mutants were kindly donated by Dr Yi-Fang Tsay. *tor-es* was kindly donated by Dr Ezequiel Petrillo, *lst8-1, rps6b, raptor1b* and overexpressing line S6K1 (S6K OE) kindly donated by Dr Elina Welchen and TOR OE and TOR OE/CA-ROP2 kindly donated by Dr. Lyubov Ryabova.

### Pharmacological Treatments

For all experiments, plants were grown first on solid 0.5X MS medium at 22°C for 5 days in continuous light. According to the specific treatment plants were transferred to plates containing regular solid 0.5X MS supplemented with 100nM IAA (auxin treatment), 250 nM AZD-8055 (TOR inhibition) or 10 μM β-estradiol (*tor-es* line low temperature treatment) and grown at 22°C for 5 days + 3 days at 10°C in continuous light. RH phenotype was quantified after each 3 days span. For nitrate response treatment plants were transferred to plates containing low or high nitrate solid media supplemented with 10 μM β-estradiol (*tor-es* line), grown at 22°C for 3 days in continuous light and RH phenotype quantified.

### Root hair phenotype

Seeds were surface sterilized and stratified in darkness for 3 days at 4°C. Then grown *in vitro* on a specific condition and medium in a plant growth chamber in continuous light (120 μmol.sec^-1^.m^-2^) at 22°C and/or 10°C. The quantitative analyses of RH phenotypes of Col o and transgenic lines were made the last day of the growth conditions described in the two previous sections. In the case of low temperature treatment, the measurements were done after 5 days at 22°C and after 3 days at 10°C. For that purpose, 10 fully elongated RH from the maturation zone were measured per root under the same conditions from each treatment and control. Images were captured using an Olympus SZX7 Zoom Stereo Microscope (Olympus, Japan) equipped with a Q-Colors digital camera and Q Capture Pro 7 software (Olympus, Japan). Results were expressed as the mean ± SD using the GraphPad Prism 8.0.1 (USA) statistical analysis software. Results are representative of three independent experiments, each involving 7–20 roots.

### Confocal Microscopy

For measurements of fluorescence intensity distributions after cold stress (22°C→10°C) in root tips and root hairs of pFER::FER-GFP and pROP2::ROP2-mCitrine lines confocal laser scanning microscopy Zeiss LSM 710 (Carl Zeiss, Germany) was used. For image acquisition, 10x/0.3 NA EC Plan-Neofluar objective for root tips, and 40x/1.4 Oil DIC Plan Apochromat objective for root hairs were used. GFP signal was excited with 488 nm argon laser at 4% laser power intensity and emission band of 493-549 nm. mCytrine signal was excited with 514 nm argon laser at 4% laser power intensity and emission band 519-583 nm. Fluorescence intensity measurements (in A.U.) were generated using Zen Black 2011 software (Carl Zeiss, Germany) and graphically edited in Microsoft Excel. Results are representative of three independent experiments, each involving 1–2 roots and 1 to 4 hairs per root.

### Co-IP assay

For Co-IP assays using A/G agarose and an anti-TOR antibody^32^, 30 μL of A/G beads (Thermo Fisher Scientific Inc., 20421) was resuspended and washed three times using NEB buffer (20 mM HEPES [pH 7.5], 40 mM KCl, 5 mM MgCl_2_) before adding 8 μL of anti-TOR antibody or preimmune serum as a negative control^32^ in a total volume of 500 μL of NEB buffer, followed by incubation for 4 h at 4 °C. Col-0 seedlings were first grown at 22 °C for 5 days then transferred to 22 °C or 10 °C for 2 days or 3 days. For protein extraction from plants, the collected materials were ground to a fine powder in liquid nitrogen and solubilized with NEB-T buffer (20 mM HEPES [pH 7.5], 40 mM KCl, 5 mM MgCl_2_, 0.5% Triton X-100) containing 1 x protease inhibitor cocktail (Thermo Fisher Scientific Inc., 78430) and 1 x phosphatase inhibitor (Thermo Fisher Scientific Inc., 78420) and incubated for 1 h on the ice. The extracts were centrifuged at 16,000 g at 4 °C for 15 min, and the resultant supernatant was incubated with prepared antibody-beads from the above step. After overnight incubation at 4°C with rotation, the agarose beads were washed five times with the NEB buffer and eluted with elution buffer (0.2 M glycine, 0.5% Triton X-100, pH 7.5). Anti-FER and anti-TOR antibodies^30^ were used for immunoblotting to detect the immunoprecipitates.

### TOR immunolocalization

Col-0 and *fer-4* seedlings were first grown at 22 °C for 5 days before transferred to 22 °C or 10 °C for 3 days. Then seedlings were collected and incubated for 10 min under vacuum (0.05 MPa) in phosphate-buffered saline (PBS) containing 4% paraformaldehyde and 0.1% Triton X-100. Seedlings were washed gently three times (10 min for each wash) in PBS and then the cell wall was digested in 2% Driselase (Sigma, D8037) in PBS for 18 min at 37 °C and washed five times with PBS. The permeability of the seedlings was increased by incubating them in 3% IGEPAL CA-630 (Sigma, 18896) and 10% DMSO in PBS for 18 min, followed by washed three times with PBS. Seedlings were incubated in 2% bovine serum albumin (BSA) (Ameresco, 0332) in PBS for 1.5 h and then incubated with primary TOR antibody3^2^ (antibody diluted 1:600 in 2% BSA) for overnight at 4°C. The seedlings were washed with PBS for five times. Fluorophore-labeled secondary antibody (goat–mouse secondary antibody, diluted 1:600 in 2% BSA) was incubated with the samples at 37°C for 5 h in the dark. Seedlings were washed five times with PBS before observation. Fluorescent signal detection and documentation were performed using a Nikon confocal laser scanning microscope with a 560-nm band-pass filter for IF555 detection.

### S6K-p/S6K immunoblotting detection

For immunological detection of S6K-p and S6K1/2, total soluble proteins were extracted from 50 mg of plant materials grown as indicated previously with 100 μL 2 x Laemmli buffer supplemented with 1% Phosphatase Inhibitor Cocktail 2 (Thermo Fisher Scientific Inc., 78430). Proteins were denatured for 10 min at 95°C and separated on 10% or 8% SDS-PAGE. Rabbit AtTOR polyclonal antibodies (Abiocode, R2854-2), rabbit polyclonal S6K1/2 antibodies (Agrisera, AS121855), and S6K1-p (phospho T449) antibody (Abcam, ab207399) and ACTIN antibody (Abmart, M20009) were used for immunoblotting.

### Quantitative PCR (qPCR)

Total root RNA was extracted from plantlets grown *in vitro* at 22 °C and 10 °C using the RNeasy^®^Plant Mini Kit (QIAGEN, Germany). One microgram of total RNA was reverse transcribed using an oligo(dT)_20_ primer and the Super Script™ IV RT (Invitrogen,USA) according to the manufacturer’s instructions. cDNA was diluted 20-fold before PCR. qPCR was performed on a LightCycler^®^480 Instrument II (Roche, USA) using 2 μL of 5X HOT FIREPol^®^EvaGreen^®^ qPCR Mix Plus(no ROX) (Solis BioDyne, Estonia), 2 μL of cDNA, and 0.25 μM of each primer in a total volume of 10 μL per reaction. ACT2 (AT3G18780) gene was used as reference for normalization of gene expression levels (ACT2 primers, primers, F: GGTAACATTGTGCTCAGTGGTGG R: CTCGGCCTTGGAGATCCACATC; TOR primers, F: GAAGATGAAGATCCCGCTGA R: GCATCTCCAAGCATATTTACAGC^44^). The cycling conditions were: 95 °C for 12 min., 35 cycles of 95°C for 15 sec., 60°C for 1 min and finally a melting curve from 60°C to 95 °C (0.05°/sec). Data were analyzed using the ΔΔC_t_ method^92^ and LightCycler^®^480 Software, version 1.5 (Roche). Two independent experiments with three biological and three technical replicates per experiment, were performed.

### Qualitative determination of KNO_3_ diffusion at 10°C and 22°C

This method was adapted from a previous protocol^93^. First, 25 ml of 0.8% agar-water media was prepared into plastic Petri dishes. After gelation it was removed the center of the agar plate by using the rear part of a p1000 tip as a puncher (removing a volume of approx. 200 μl of gel). After equilibration at either 10°C or 22°C it was added 200 μl of liquid 0.8% Agar (at 40°C) containing 0,94M KNO_3_ to fill up the well and started the diffusion time. After 3 h of incubation, 20 μl agar disks were taken at different distances from the border of the KNO_3_ zone (from 5mm to 25mm in 5mm steps). For this purpose, we used a Pasteur pipette as a puncher. The agar disks were incubated with 200 μl of 2mM diphenylamine ((C_6_H_5_)2NH) colorless solution in 14,4M H_2_SO_4_ with eventual shaking to homogenize the reaction media. After 30 min we transferred 100ul of the bright blue resultant reaction to a 96 well plate and measured OD at 595 nm. Serial dilutions showed deviations from the expected linear behavior preventing the possibility of a robust quantitative assay. We used high KNO_3_ concentrations to prevent excessive dilution of the NO_3_^-^ below detection levels and speed up the diffusion process.

## Acknowledgements

We would like to thank Elina Welchen, Ezequiel Petrillo, Yi-Fang Tsay, Kriss Vissenberg for the seed lines. We thank NASC (Ohio State University) for providing T-DNA lines seed lines. J.M.E. is investigator of the National Research Council (CONICET) from Argentina. M.I. is supported by ANID FONDECYT POSTDOCTORADO [grant 3220138]. This work was supported by grants Natural Science Foundation of China (NSFC-32001974 and NSFC-31871396) to F.Y. and (NSFC-31900232) to L.S. and from ANPCyT (PICT2017-0066, and PICT2019-0015), by ANID - Programa lniciativa Cientifica Milenio ICN17_022, NCN2021_010 and Fonda Nacional de Desarrollo Cientifico y Tecnol6gico [1200010] to J.M.E., by the Deutsche Forschungsgemeinschaft (DFG; GR4559_ 4-1, GR4559_5-1, EXC-2048/1 project ID 390686111) to G.G., and by the Czech Science Foundation GACR (project Nr. 19-18675S) to J. Š.

## Author Contribution

J.M.P performed most of the experiments, analysed the data and helped in the writing process of the manuscript. LS. performed the S6K-p determinations, IP of TOR-FER and immunolocalization assay of TOR. L.K., M.O. and J.Š. provided live cell imaging data on FER and ROP2. V.B.G., J.M.P., T.U.I., M.A.I., M.M.R., S-Z., Y.S., R.A.G., M.S., L.A.R., J.M.A., G.G., J. Š. helped in the data analysis and writing process of the manuscript. F.Y. designed research and analysed part of the data and J.M.E. designed research, analysed the data, supervised the project, and wrote the paper. All authors commented on the results and the manuscript. This manuscript has not been published and is not under consideration for publication elsewhere. All the authors have read the manuscript and have approved this submission.

