## Supplementary Files for "Cell surface receptor kinase FERONIA linked to nutrient sensor TORC signaling controls root hair growth at low temperature linked to low nitrate in *Arabidopsis thaliana*"

Javier Martinez Pacheco et al.

**This File includes:**

Supplementary Tables 1-2

Supplementary Figures 1-8

**Table S1**. Recipe of nutritional media used in this study. MS 0.5X including MES from Duchefa (https://www.duchefa-biochemie.com/product/details/number/M0254/name/murashige-skoog-medium-incl-mes-buffer). Commercial media M407 (lacking KNO_3_, NH_4_NO_3_, or KH_2_PO_4_) (https://phytotechlab.com/murashige-skoog-modified-basal-salt-mixture-m407.html).

| **Nutrients** | **Commercial MS 0.5X (mg/l)** | **Commercial M407 0.5X (mg/l)** | **M407 0.5X low *vs* high N experiments (mg/l)** |
| --- | --- | --- | --- |
| KNO_3_ | 950 | -- | 50 (low N, equivalent to MS 0.025X) or 1900 (high N, equivalent to MS 1X) |
| NH_4_NO_3_ | 825 | -- | -- |
| KH_2_PO_4_ | 85 | -- | 85 |
| CaCl_2_ anhydrous | 166 | 166 | 166 |
| MgSO_4_ anhydrous | 90.2 | 90.2 | 90.2 |
| CoCl_2_.6H_2_O | 0.0125 | 0.0125 | 0.0125 |
| CuSO_4_.5H_2_O | 0.0125 | 0.0125 | 0.0125 |
| FeNaEDTA | 18.35 | -- | -- |
| 2NaEDTA.2H_2_O | -- | 18.63 | 18.63 |
| FeSO_4_.7H_2_O | -- | 13.9 | 13.9 |
| H_3_BO_3_ | 3.1 | 3.1 | 3.1 |
| KI | 0.415 | 0.415 | 0.415 |
| MnSO_4_.H_2_O | 8.45 | 8.45 | 8.45 |
| Na_2_MoO_4_.2H_2_O | 0.125 | 0.25 | 0.125 |
| ZnSO_4_.7H_2_O | 4.3 | 4.3 | 4.3 |
| MES Buffer | 250 | -- | 250 |
| Working Solution pH | 5.75 | 5.75 | 5.75 |
| Plant Agar (g/l) | 8 | 8 | 8 |

**Table S2. List of mutant and transgenic lines used in this study.**

| **Name** | **Reference** |
| --- | --- |
| *fer-4* | Duan *et al*., 2010 |
| *fer-5* |  |
| *fer-8* | Song *et al*., 2021 |
| FER K565R | Escobar-Restrepo *et al*., 2007 |
| *p*FER:FER-GFP |  |
| *eru* | Schoenaers *et al*., 2018 |
| TOR RNA*i* estradiol inducible (*tor-es*) | Xiong and Sheen, 2012 |
| *lst8-1* | Moreau *et al*., 2012 |
| *raptor 1b* | Deprost *et al*., 2005 |
| *rps6b* | Creff *et al*., 2010 |
| S6K OE | Schepetilnikov *et al*., 2017 |
| 35S:TOR-GFP (TOR OE) | Schepetilnikov *et al*., 2017 |
| 35S:TOR-GFP/CA ROP2 (TOR OE/CA ROP2) |  |
| *chl1-5* | Ho *et al*., 2009 |
| *chl1-9* |  |
| NRT1.1 T101D |  |
| NRT1.1 T101A |  |
| *rop2* | Denninger *et al*., 2019 |
| *rop2 rop4* |  |
| *p*ROP2:ROP2-mCitrine |  |
| ROP2 OE/Col-0 | Song *et al*., 2021 |
| ROP2 OE/*fer-8* #1-11 |  |
| ROP2 OE/*fer-8* #2-7 |  |
| ROP2 OE/*fer-8* #3-1 |  |
| *tga1 tga4* | Kesarwani *et al*., 2007 |
| 35S:TGA1 | Switf et al. 2020 |

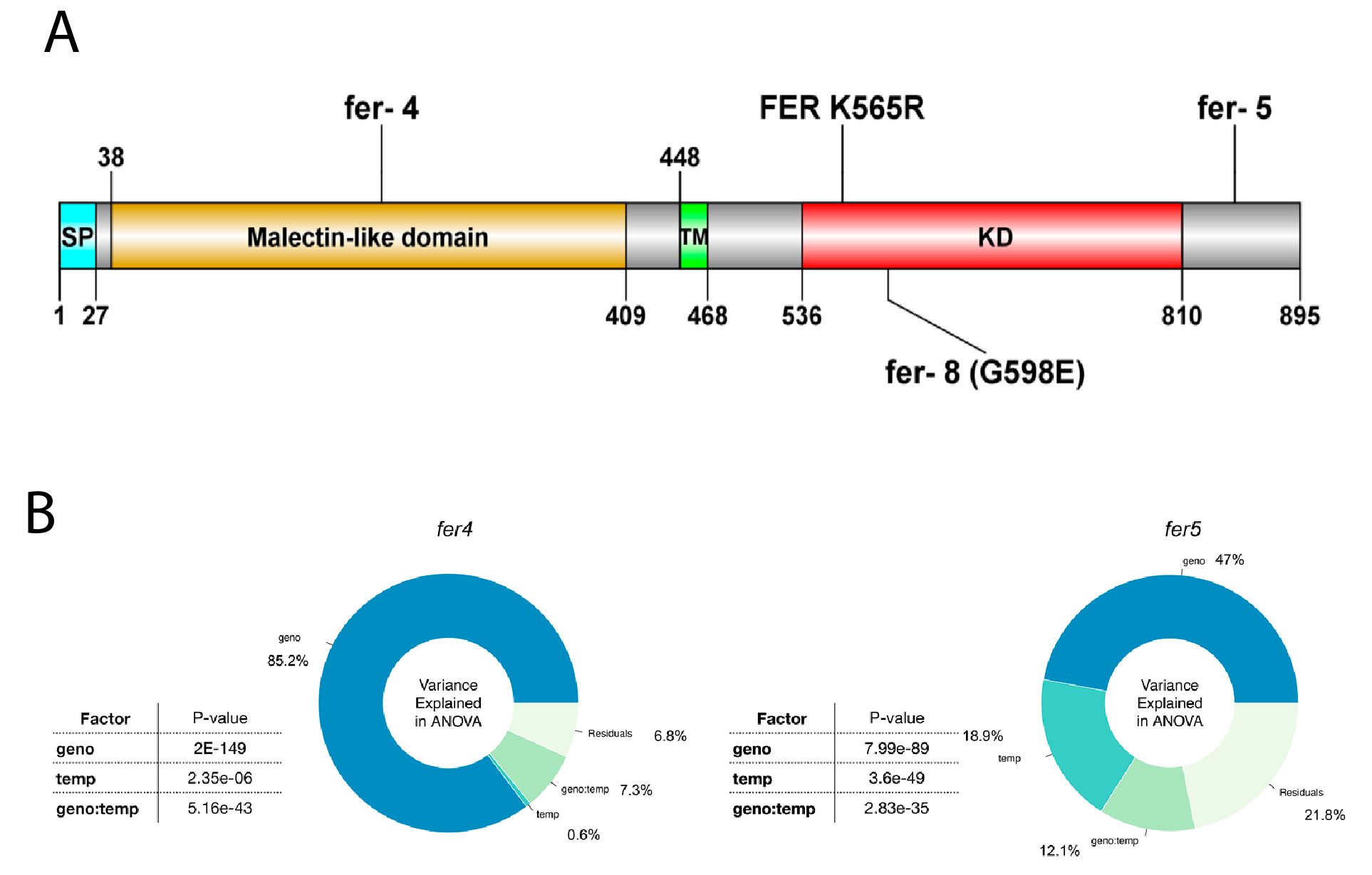

**Figure S1A. FER protein domains and insertion/mutation positions of the alleles described in this study.** Diagram created using DOG software v. 2.0. **Figure S1B**. FER affect general RH growth and elongation of RH in response to cold. The contribution (percentage %) of each factor in a two-way ANOVA model was represented in a pie chart. The p-value of the ANOVA analysis of the factors: genotype (Geno), temperature (Temp), and the interaction between genotype and temperature (Geno:Temp) is indicated for each comparison between Col-0 and mutant/altered gene.

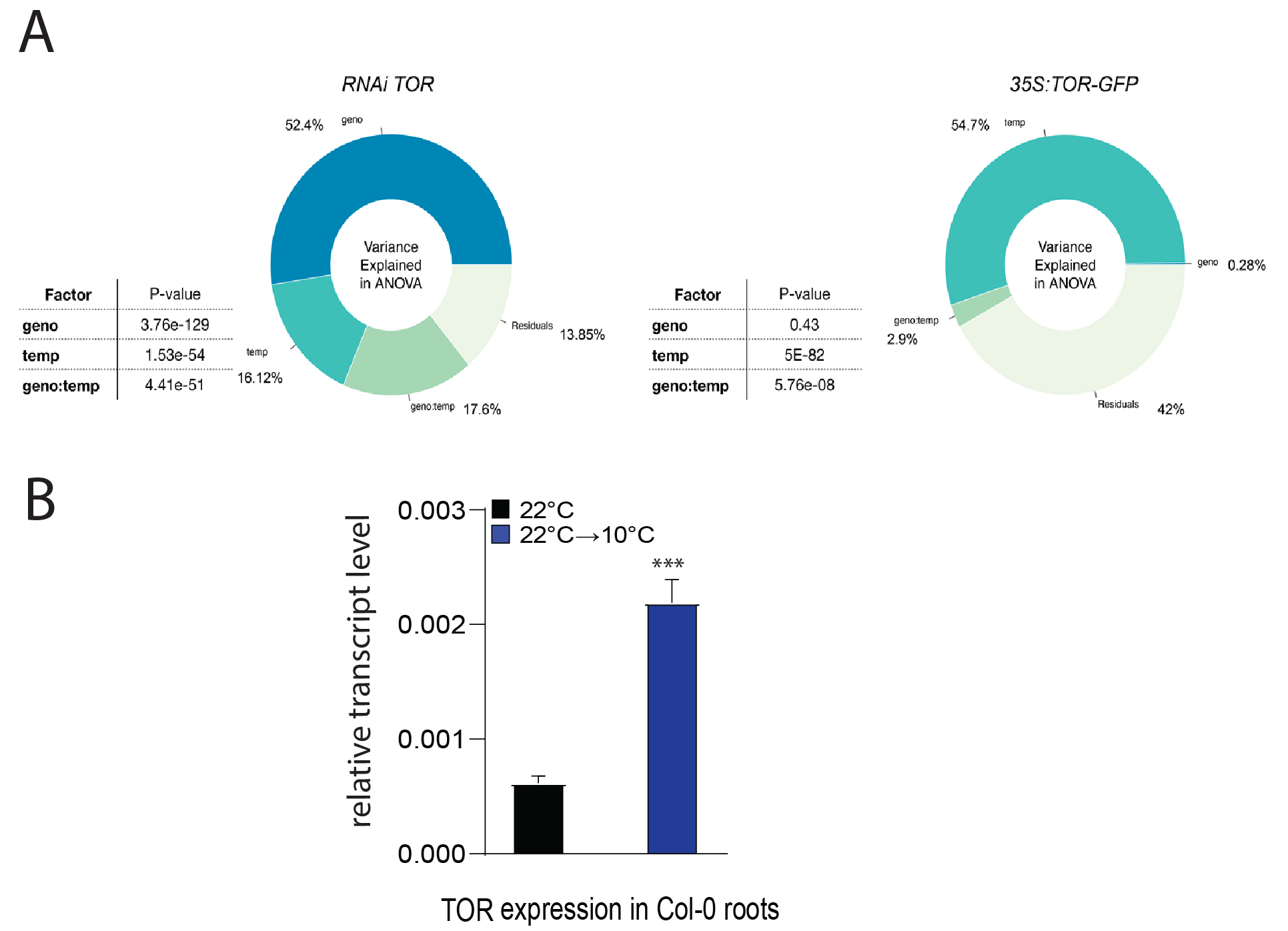

**Figure S2A. TOR affects general RH growth and elongation of RH in response to cold.** The contribution (percentage %) of each factor in a two-way ANOVA model was represented in a pie chart. The p-value of the ANOVA analysis of the factors: genotype (Geno), temperature (Temp), and the interaction between genotype and temperature (Geno:Temp)) is indicated for each comparison between Col-0 and mutant/altered gene. **S2B.** Quantitative PCR of *TOR* expression levels in Col-0 roots grown at 22^o^C and 10^o^C. *ACT2* expression was used for normalization of gene expression. Three biological replicates and three technical replicates per experiment were performed**.** Asterisks (***) indicate significant differences according to unpaired *t* test, two-tailed *p*<0.05.

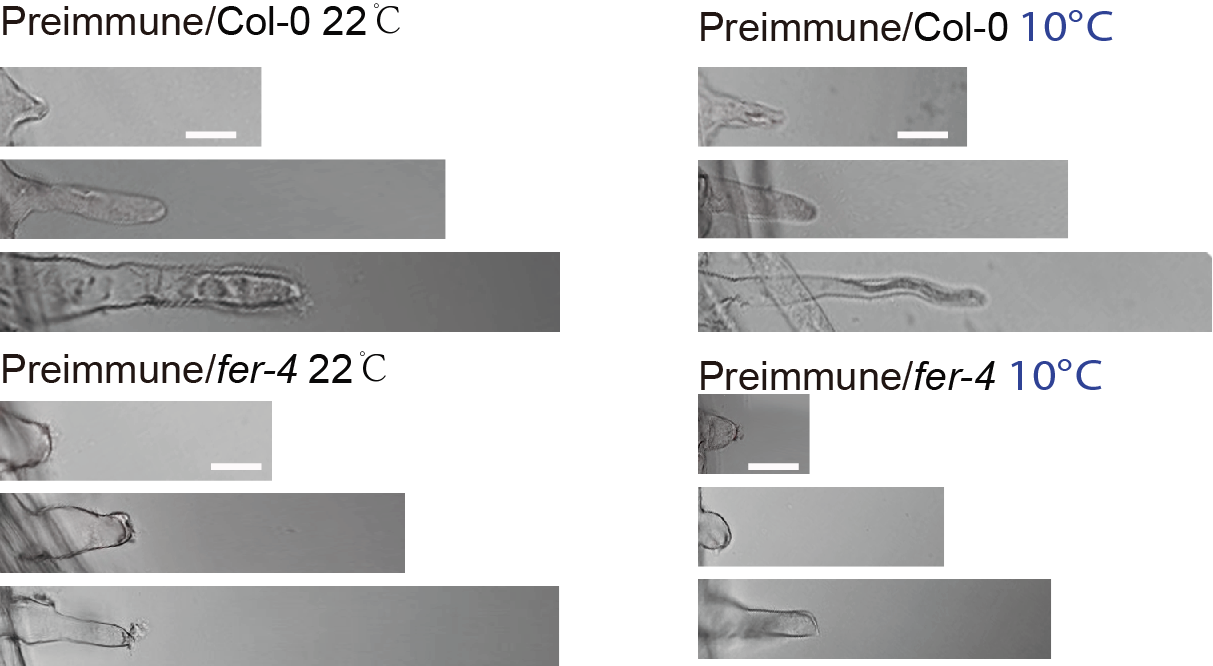

**Figure S3**. Representative control images of the TOR immunolocalization experiment in RHs using preimmune serum as primary antibody. Scale bar=10 µm.

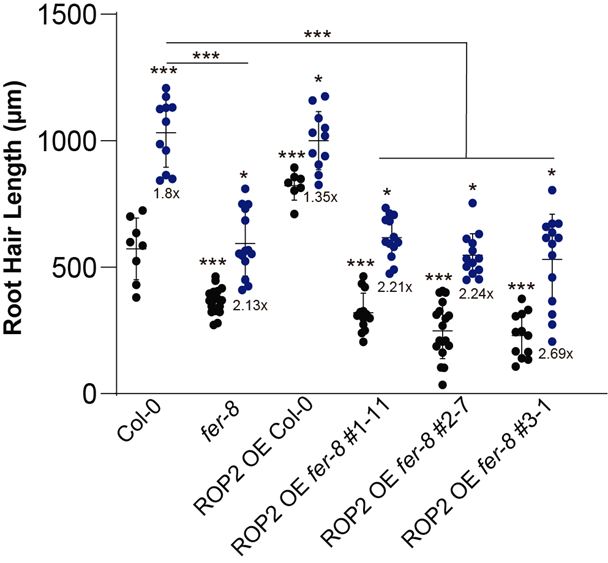

**Figure S4. ROP2 require FER to trigger RH growth.** Each point is the mean of the length of the 10 longest RHs identified in a single root. Data are the mean ± SD (N=7-20 roots), two-way ANOVA followed by a Tukey–Kramer test; (*) *p*<0.05, (***) *p*<0.001. Results are representative of three independent experiments. Asterisks indicate significant differences. Numbers under the plots represents RH growth ratio 10^o^C/22^o^C.

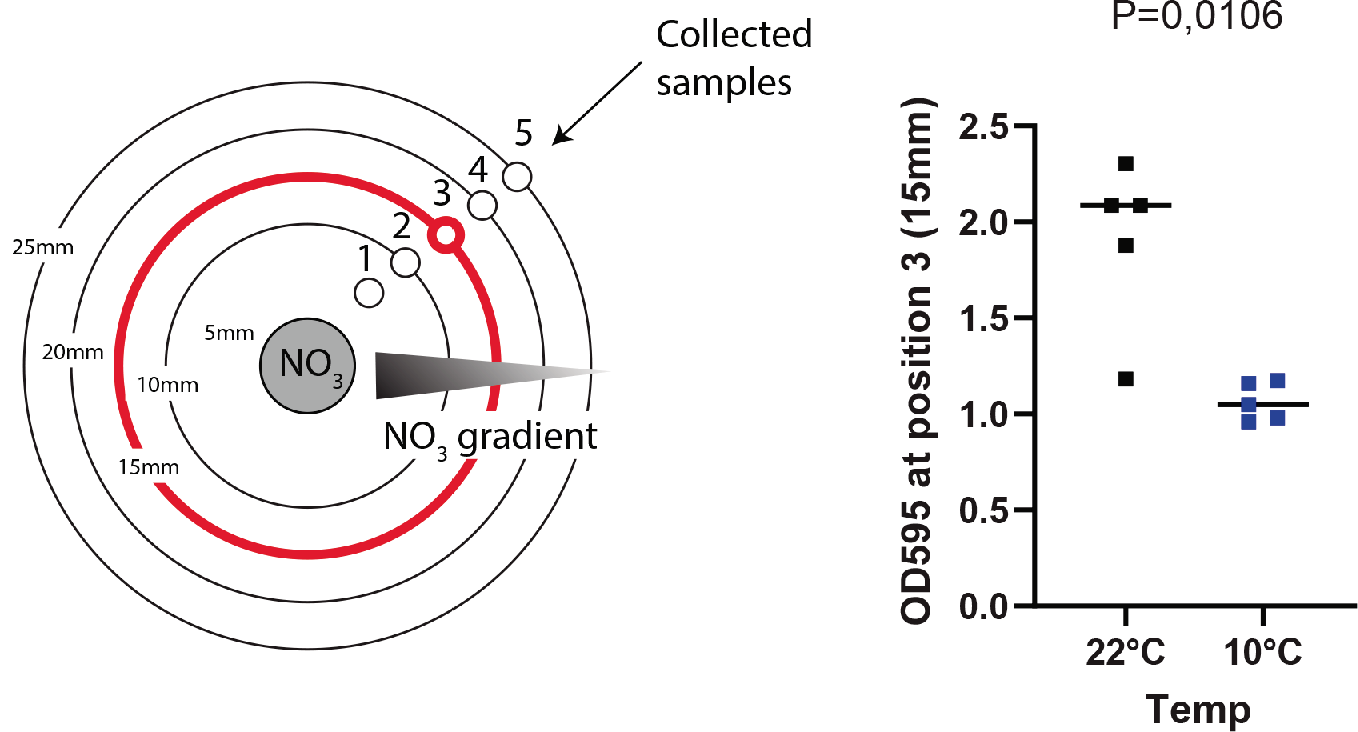

**Figure. S5. Temperature effect on the diffusion of nitrates at 22^o^C and 10^o^C**. A well was carved in the center of 0.8% water-agar plates and fi­lled with 0,94M KNO_3_ solution containing agar 0.8%. After 3h of incubation, the plate was probed by removing agar cylinders at every 5mm from the loaded center (up to 25 mm). The cylinders were directly incubated in diphenylamine solution in 14,4M H_2_SO_4_ and incubated for 30 min. OD_595_ was then determined. Only samples taken from the 3rd line (15mm) showed non-saturated and detectable OD values. Significant differences according to unpaired *t* test, two-tailed *p*<0.05.

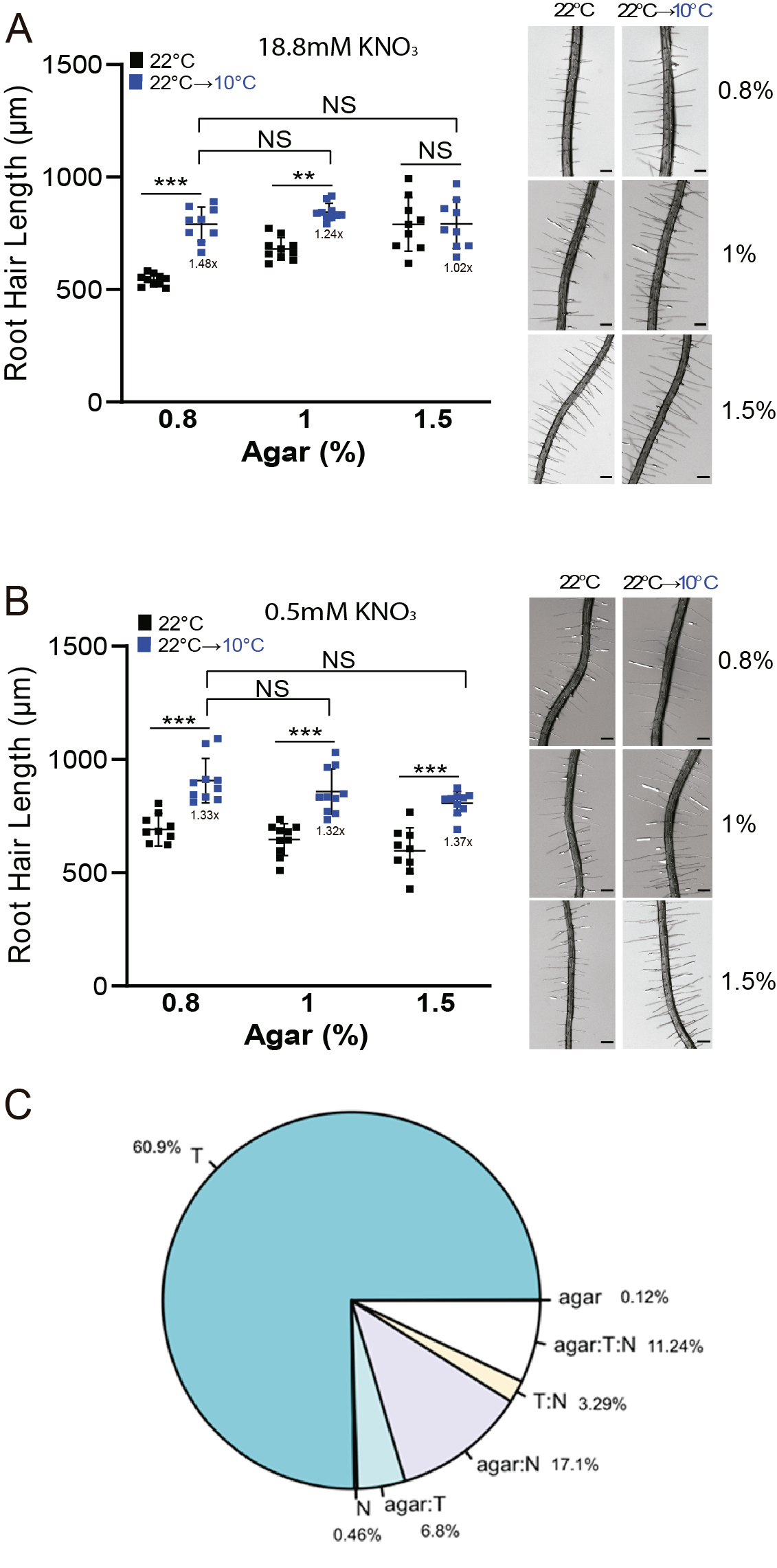

**Figure S6. Lower mobility of nitrate induced by higher agar concentration (1,5%) regulates the RH growth independent of the temperature. (A)** High nitrogen and **(B)** Low Nitrogen. Each point is the mean of the length of the 10 longest RHs identified in a single root. Data are the mean ± SD (N= 10 roots), two-way ANOVA followed by a Tukey–Kramer test; (*) *p*<0.05, (**) *p*<0.01, (***) *p*<0.001. Results are representative of three independent experiments. Asterisks indicate significant differences. Representative images of each line are shown on the right. Scale bars=300 µm. Numbers under the plots represents RH growth ratio 10^o^C/22^o^C. (**C**) The contribution (percentage %) of each factor in a three-way ANOVA model was represented in a pie chart. Three-way ANOVA analysis across all conditions (Figure S4A and S4B) reveals that temperature (T), the interaction of agar with NO3- (agar:N); and the interaction of agar, temperature, and NO3- (agar:T:N) largely explain the variance of RH elongation; while NO3- and agar factors on their own are less influential.

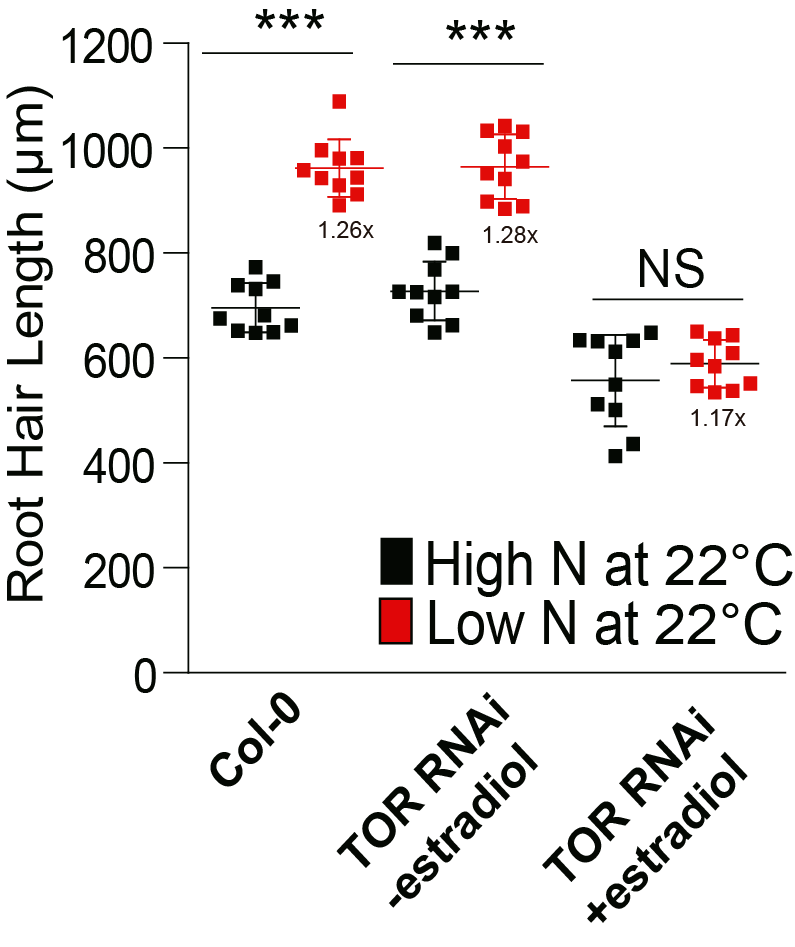

**Figure S7**. **Low nitrate perception relays in TOR to trigger RH growth.** Scatterplot of RH length of Col-0 and *tor-es* line grown in low and high N conditions. Differential growth of RH at low N is suppressed in the estradiol inducible *RNAi TOR* line. Each point is the mean of the length of the 10 longest RHs identified in the maturation zone of a single root. Data are the mean ± SD (N=7-10 roots), two-way ANOVA followed by a Tukey–Kramer test; (***) *p*<0.001, NS=non-significant. Results are representative of three independent experiments. Asterisks indicate significant differences between the same genotype at different N concentration or between different genotypes at the same N concentration. Numbers under the plots represents RH growth ratio low N/high N.

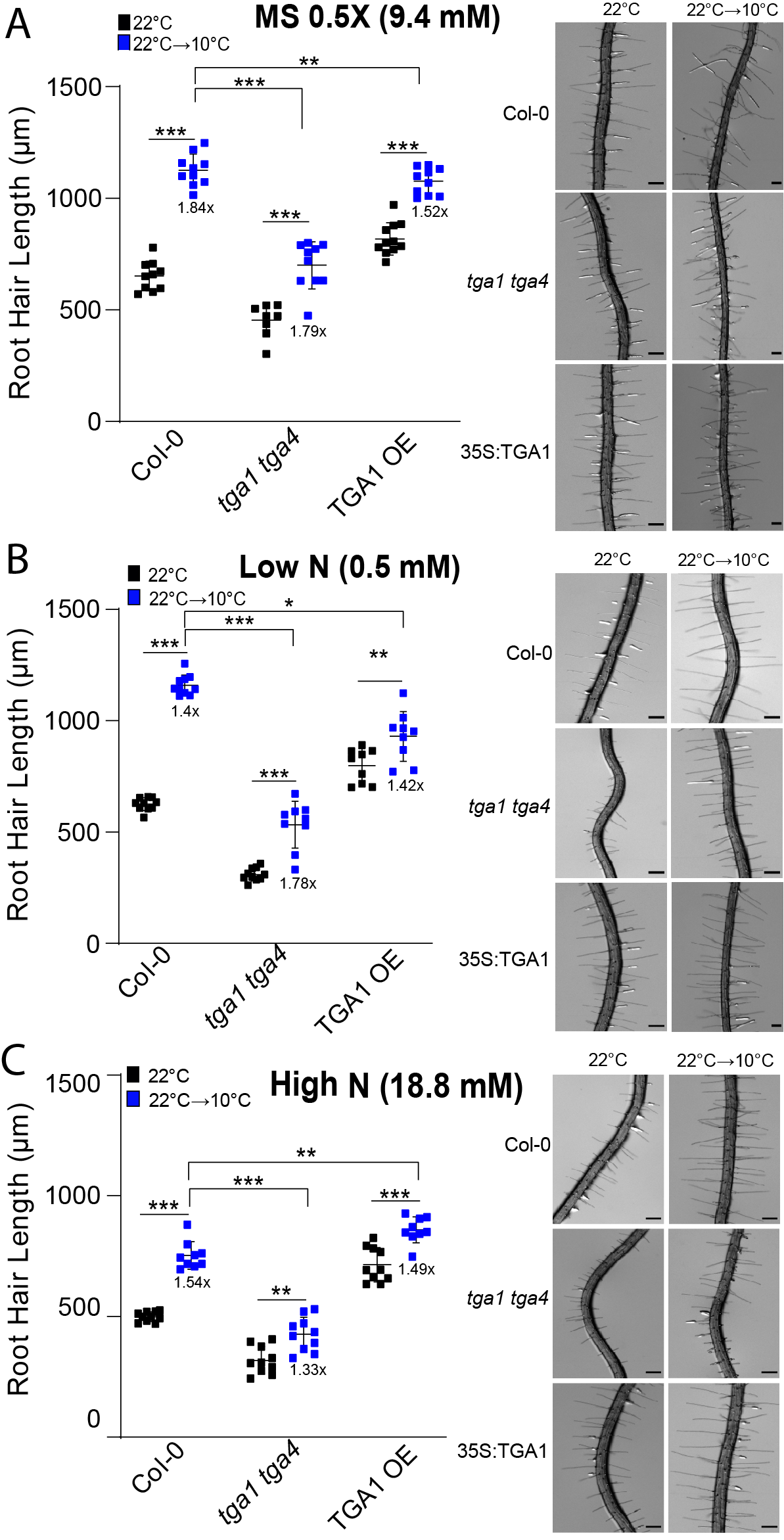

**Figure S8. TGA1 TG4 regulates the RH growth under low temperature.**

**(A)** MS 0.5X (9.4mM nitrates), (**B**) Low N (nitrogen, 0.5 mM) and **(B**) High N (Nitrogen, 18.8mM). Each point is the mean of the length of the 10 longest RHs identified in a single root. Data are the mean ± SD (N= 10 roots), two-way ANOVA followed by a Tukey–Kramer test; (*) *p*<0.05, (**) *p*<0.01, (***) *p*<0.001. Results are representative of three independent experiments. Asterisks indicate significant differences. Representative images of each line are shown on the right. Scale bars=300 µm. Numbers under the plots represents RH growth ratio 10^o^C/22^o^C.
